# Senataxin and DNA-PKcs Redundantly Promote Non-Homologous End Joining Repair of DNA Double Strand Breaks During V(D)J Recombination

**DOI:** 10.1101/2024.09.25.615014

**Authors:** Bo-Ruei Chen, Thu Pham, Lance D. Reynolds, Nghi Dang, Yanfeng Zhang, Kimberly Manalang, Gabriel Matos-Rodrigues, Jason Romero Neidigk, Andre Nussenzweig, Jessica K. Tyler, Barry P. Sleckman

## Abstract

Non-homologous end joining (NHEJ) is required for repairing DNA double strand breaks (DSBs) generated by the RAG endonuclease during lymphocyte antigen receptor gene assembly by V(D)J recombination. The Ataxia telangiectasia mutated (ATM) and DNA-dependent protein kinase catalytic subunit (DNA-PKcs) kinases regulate functionally redundant pathways required for NHEJ. Here we report that loss of the senataxin helicase leads to a significant defect in RAG DSB repair upon inactivation of DNA-PKcs. The NHEJ function of senataxin is redundant with the RECQL5 helicase and the HLTF translocase and is epistatic with ATM. Co-inactivation of ATM, RECQL5 and HLTF results in an NHEJ defect similar to that from the combined deficiency of DNA-PKcs and senataxin or losing senataxin, RECQL5 and HLTF. These data suggest that ATM and DNA-PKcs regulate the functions of senataxin and RECQL5/HLTF, respectively to provide redundant support for NHEJ.

## Introduction

The genes that encode antigen receptors are assembled in developing lymphocytes through the process of V(D)J recombination (*1, 2*). V(D)J recombination is initiated when the RAG1 and RAG2 proteins, which together form the RAG endonuclease, introduce DNA double strand breaks (DSBs) at the borders of a pair of recombining variable (V), diversity (D) or joining (J) gene segments and their adjacent RAG recognition sequences termed recombination signal sequences (RSSs) (*3, 4*). This reaction leads to the formation of a blunt signal end and a hairpin-sealed coding end at each RAG DSB. These DNA ends are then processed and repaired by the non-homologous end joining (NHEJ) pathway of DNA DSB repair. The two blunt signal ends are directly ligated to generate a signal join while the two hairpin-sealed coding ends are first opened by Artemis before ligation to form a coding join (*5, 6*). Signal ends join precisely with occasional loss of a few nucleotides whereas coding ends are joined imprecisely with most joins undergoing nucleotide gain or loss (*5, 6*).

The NHEJ pathway of DSB repair functions in all phases of the cell cycle, with the exception of mitosis, and relies on the core factors KU70, KU80, DNA Ligase IV and XRCC4, which are all essential for NHEJ-mediated DSB repair (*7-10*). Loss of any of these factors leads to a complete block in NHEJ-mediated DSB repair. Additional NHEJ factors are required for the repair of DSBs with unique end structures, such as Artemis, a nuclease that is needed to open the hairpin sealed coding ends during V(D)J recombination (*11*). In addition, accessory proteins, including 53BP1, H2AX, XLF, PAXX and MRI, promote efficient NHEJ with the individual loss of any of these proteins leading to a modest defect in NHEJ while the combined inactivation of some of these factors severely impairs NHEJ-mediated DSB repair (*12*).

The serine/threonine kinases Ataxia telangiectasia mutated (ATM) and DNA-dependent protein kinase catalytic subunit (DNA-PKcs) are activated early in the response to DNA DSBs to regulate pathways required for NHEJ via phosphorylation of many downstream effectors (*7, 13*). DNA-PKcs is activated by DSBs upon recruitment to broken DNA ends by the KU70/KU80 heterodimer, while ATM activation in response to DSBs is mediated by the MRE11/RAD50/NBS1 (MRN) complex (*14-17*). Although ATM and DNA-PKcs are known to mediate many important processes during the DNA damage response and repair, loss of either ATM or DNA-PKcs only leads to limited defects in NHEJ. In contrast, the combined loss of function of both kinases results in a stronger defect in NHEJ demonstrating that they control key NHEJ repair activities that function redundantly (*18-21*).

To reveal novel factors that are regulated by ATM or DNA-PKcs kinases and function redundantly during NHEJ, we carried out a CRISPR/Cas9 whole genome guide RNA (gRNA) screen in Abelson murine leukemia viral kinase-transformed pre-B cell lines, hereafter referred to as abl pre-B cells. Treatment of abl pre-B cells with the abl kinase inhibitor imatinib leads to arrest in G_0_ phase and initiation of V(D)J recombination (*22-24*). We found that gRNAs to the senataxin gene (*Setx*) strongly impaired NHEJ-mediated repair of RAG DSBs in G_0_-arrested abl pre-B cells treated DNA-PKcs inhibitor NU7441, but not in the untreated cells. Senataxin is a member of the superfamily 1 (SF1) RNA helicases that unwinds RNA/DNA hybrids, including R-loops, a three-strand structure composed of an RNA/DNA hybrid and a displaced non-template single strand DNA (ssDNA) (*25-28*). Here we establish that senataxin is required for NHEJ in cells where DNA-PKcs is inactivated and that senataxin acts redundantly with the RECQL5 helicase and the HLTF translocase. This is the first evidence implicating multiple collaborating helicase and translocase activities in NHEJ in ATM- and DNA-PKcs-dependent manner.

## Results

### Senataxin is required for RAG DSB repair in DNA-PKcs-deficient abl pre-B cells

V(D)J recombination in abl pre-B cells can be assayed with chromosomally integrated retroviral recombination substrates that contain a single pair of recombination signal sequences (RSSs) (*23, 29*). The pMG-INV recombination substrate contains an anti-sense GFP cDNA flanked by RSSs oriented such that successful V(D)J recombination leads to inversion of the GFP cDNA and GFP expression (Fig. 1A) (*29*). To identify genes critical for NHEJ-mediated RAG DSB repair in cells treated with a DNA-PKcs kinase inhibitor, wild type (WT) abl pre-B cells containing the pMG-INV recombination substrate were transduced with a whole genome lentiviral gRNA library and treated with imatinib to induce V(D)J recombination in the presence of the DNA-PKcs inhibitor NU7441 or vehicle alone. Cells that completed V(D)J recombination (GFP^+^) and those that did not (GFP^-^) were isolated by fluorescence activated cell sorting (FACS) and the gRNAs in these cells were amplified by PCR and sequenced (Fig. 1B). Among the gRNAs highly enriched in GFP^-^ populations were those to the core NHEJ factors and genes required for generating RAG DSBs, such as *Lig4* and *Rag1*, respectively, thus validating our screening approach (Table S1). The gRNAs to *Setx*, a gene which encodes the protein senataxin, were 3- to 5-fold more enriched in the GFP^-^ population of abl pre-B cells treated with NU7441 as compared to vehicle alone, suggesting that senataxin is required for V(D)J recombination in abl pre-B cells where DNA-PKcs kinase activity has been inhibited (Fig. 1C).

**Figure 1:**
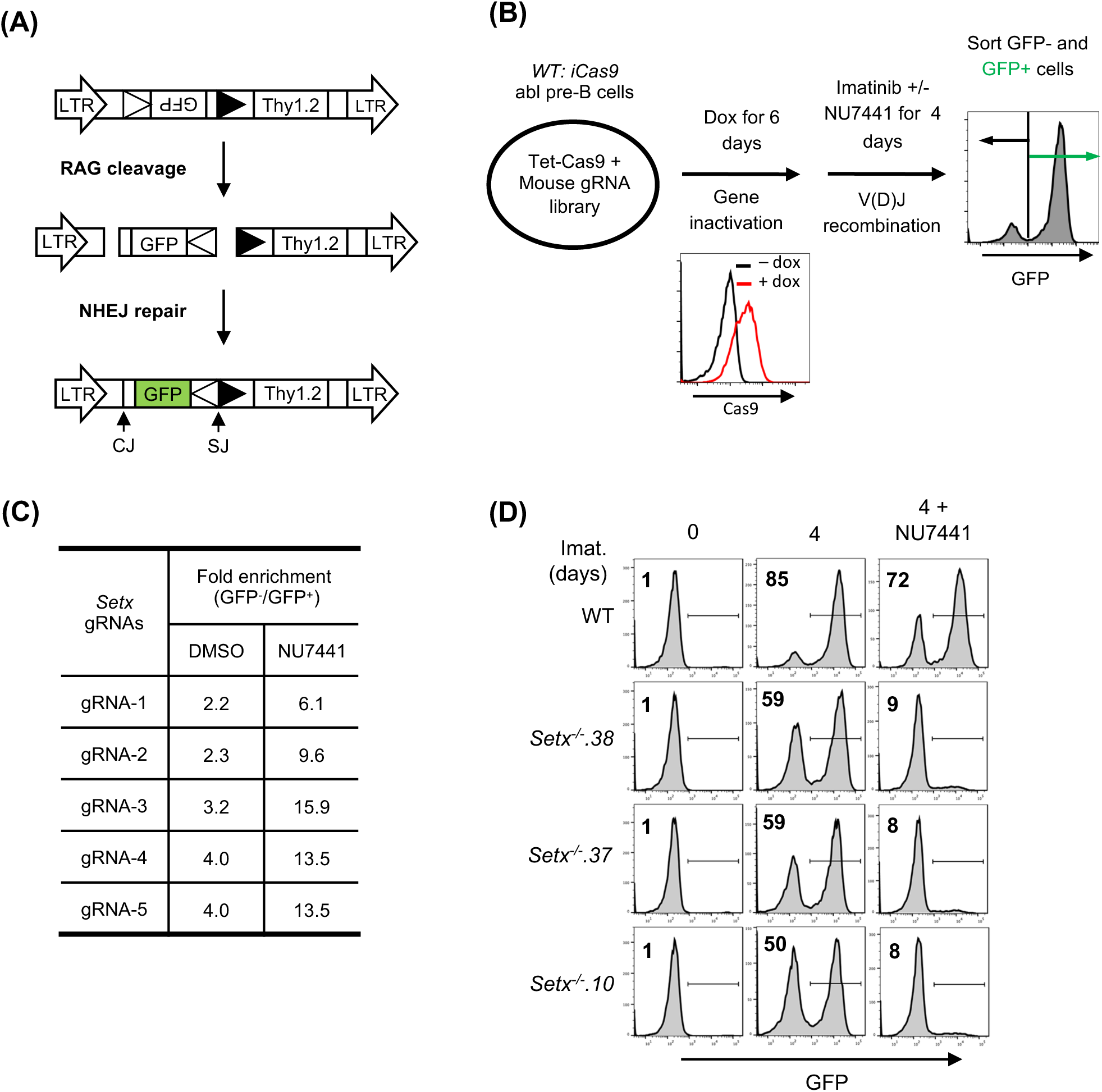
A Genome-wide CRISPR/Cas9 screen for identifying genes required for V(D)J recombination in DNA-PKcs inhibited abl pre-B cells. (A) The schematic of the retroviral V(D)J recombination substrate pMG-INV. The open and filled triangles represent the RSSs. The retroviral LTRs, CEs and SEs upon RAG cleavage as well as CJs and SJs after NHEJ-mediated repair are indicated. The anti-sense GFP cDNA is inverted to the sense orientation upon completion of the recombination and its expression driven by the retroviral LTR (green). (B) The schematic diagram of a genome-wide CRISPR/Cas9 screen in WT abl pre-B cells for identifying genes important for V(D)J recombination upon DNA-PKcs inhibition with NU7441. (C) The fold enrichments of *Setx* gRNAs from screens conducted in DMSO and NU7441-treated WT abl pre-B cells (both also treated with imatinib). The fold enrichment is calculated as the ratio of the normalized count of a gRNA from GFP^-^ cells over that from GFP^+^ cells. (D) Flow cytometric analysis for GFP expression in WT and three independently isolated *Setx*^-/-^ abl pre-B cell lines with pMG-INV and treated with imatinib (imat.) in the presence or absence of the DNA-PKcs kinase inhibitor NU7441 for the indicated times. The percentages of GFP^+^ cells are indicated in the top left corners of the histograms.

*Setx*^-/-^ abl pre-B cell clones were generated by CRISPR/Cas9 and the inactivating mutations were verified by sequencing (data not shown). Flow cytometric analyses of GFP expression revealed that in comparison to WT abl pre-B cells, *Setx*^-/-^ abl pre-B cells with integrated pMG-INV exhibited a mild reduction in V(D)J recombination efficiency as evidenced by the slightly diminished percentage of GFP^+^ cells after imatinib treatment. However, treatment with the DNA-PKcs kinase inhibitor led to a dramatic reduction in GFP^+^ *Setx*^-/-^ abl pre-B cells as compared to WT abl pre-B cells (Fig. 1D). We conclude that senataxin is required for efficient V(D)J recombination in abl pre-B cells that have compromised DNA-PKcs kinase activity.

A deficiency in V(D)J recombination could result from defected RAG cleavage and/or impaired NHEJ-mediated repair of RAG DSBs. To determine whether senataxin is required for the repair of RAG DSBs during V(D)J recombination, we analyzed WT and *Setx*^-/-^ abl pre-B cells with pMX-DEL^CJ^ or pMX-DEL^SJ^ retroviral recombination substrates that allow for visualizing the repair of structurally different coding ends (CEs) and signal ends (SEs) and the formation of coding joins (CJs) and signal joins (SJs), respectively (Figs. 2A and 2B). The recombination of pMX-DEL^CJ^ and pMX-DEL^SJ^, which have RSSs positioned in opposite orientations, is accompanied by loss of the GFP cDNA bordered by RSSs. This creates either two chromosomal hairpin sealed CEs (pMX-DEL^CJ^) or open, blunt SEs (pMX-DEL^SJ^) to be joined by NHEJ to form CJs or SJs (Figs. 2A and 2B). Southern blot analyses of *Setx*^-/-^ abl pre-B cells containing pMX-DEL^CJ^ or pMX-DEL^SJ^ treated with imatinib and NU7441 revealed a marked reduction in the formation of CJs and SJs and an increased accumulation of unrepaired CEs and SEs, respectively, as compared to imatinib treatment alone (Figs. 2C, 2D, S1A and S1B). In contrast, treatment of WT abl pre-B cells containing pMX-DEL^CJ^ or pMX-DEL^SJ^ with imatinib and NU7441 had minimal effects on CJ and SJ formation (Figs. 2C, 2D, S1A and S1B). These results suggest that combined deficiencies in DNA-PKcs and senataxin lead to a NHEJ defect during V(D)J recombination.

**Figure 2.**
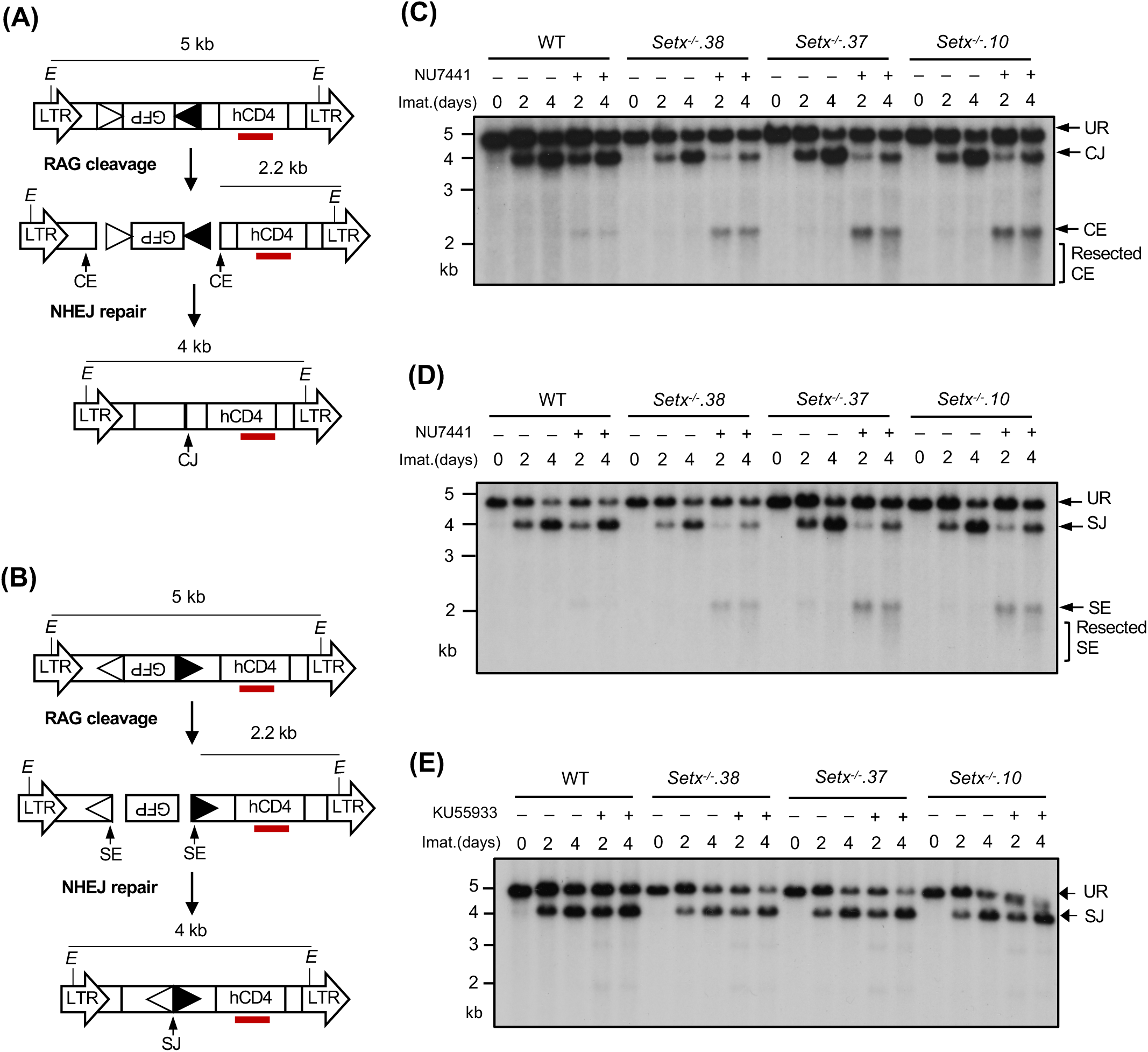
Loss of senataxin severely impairs NHEJ-mediated RAG DSB repair in DNA-PKcs-inhibited abl pre-B cells during V(D)J recombination. (A, B) The schematic of the retroviral V(D)J recombination substrates pMX-DEL^CJ^ *(A)* and pMX-DEL^SJ^ *(B)*. The open and filled triangles represent the RSSs. The red bar indicates the probe (hCD4 cDNA) for Southern blot. The Es denote the *EcoR*V restriction sequences. The retroviral LTRs, CEs and SEs upon RAG cleavage as well as CJs and SJs after NHEJ-mediated repair are indicated. (C, D) Southern blot analysis of genomic DNA from WT and three clonal *Setx*^-/-^ abl pre-B cell lines with pMX-DEL^CJ^ *(C)* or pMX-DEL^SJ^ *(D)* treated with imatinib (imat.) in the presence or absence of the DNA-PKcs kinase inhibitor NU7441 for the indicated times. The genomic DNA samples were digested with *EcoR*V and hybridized to the hCD4 probe. The restriction fragments corresponding to unrearranged reporters (UR), repaired CJs or SJs and unrepaired CEs or SEs are indicated. R. CE and R. SE indicate resected CE and SE restriction fragments. (E) Southern blot analysis of *EcoR*V digested genomic DNA from cells described in (*C*) and (*D*) treated with imatinib in the presence or absence of the ATM kinase inhibitor KU55933 for the indicated times.

We also generated *Setx^-^*^/-^: *Prkdc^-/-^*abl pre-B cells that lack both senataxin and DNA-PKcs proteins by inactivating *Prkdc*, which encodes DNA-PKcs, in *Setx*^-/-^ abl pre-B cells using CRISPR/Cas9 gene editing (Figs. 3A and S2A). Although DNA-PKcs kinase activity is not required for opening the hairpin-sealed CEs by the nuclease ARTEMIS, DNA-PKcs protein is essential for this process. Therefore, CEs remain hairpin-sealed and CJs cannot be formed in *Prkdc* null cells during V(D)J recombination (*11, 21, 30*). For this reason, only the pMX-DEL^SJ^ recombination substrate was used in this experiment. As compared to WT and *Prkdc^-/-^* abl pre-B cells, *Setx^-^*^/-^: *Prkdc^-/-^* abl pre-B cells exhibited a marked reduction in pMX-DEL^SJ^ SJ formation and accumulated unrepaired SEs after induction of V(D)J recombination (Figs. 3B and S2B). We conclude that while the loss of senataxin or DNA-PKcs kinase activity has minimal effects on RAG DSB repair, the combined loss of senataxin and DNA-PKcs kinase activity leads to a significant block in signal and coding joining during V(D)J recombination. In contrast to DNA-PKcs, inhibition of ATM kinase activity by treatment of *Setx*^-/-^ abl pre-B cells with KU55933 had no effect on SJ formation (Fig. 2E). Collectively, our results suggest that senataxin has important activities during NHEJ that are functionally redundant and uniquely shared with pathways downstream of DNA-PKcs.

**Figure 3:**
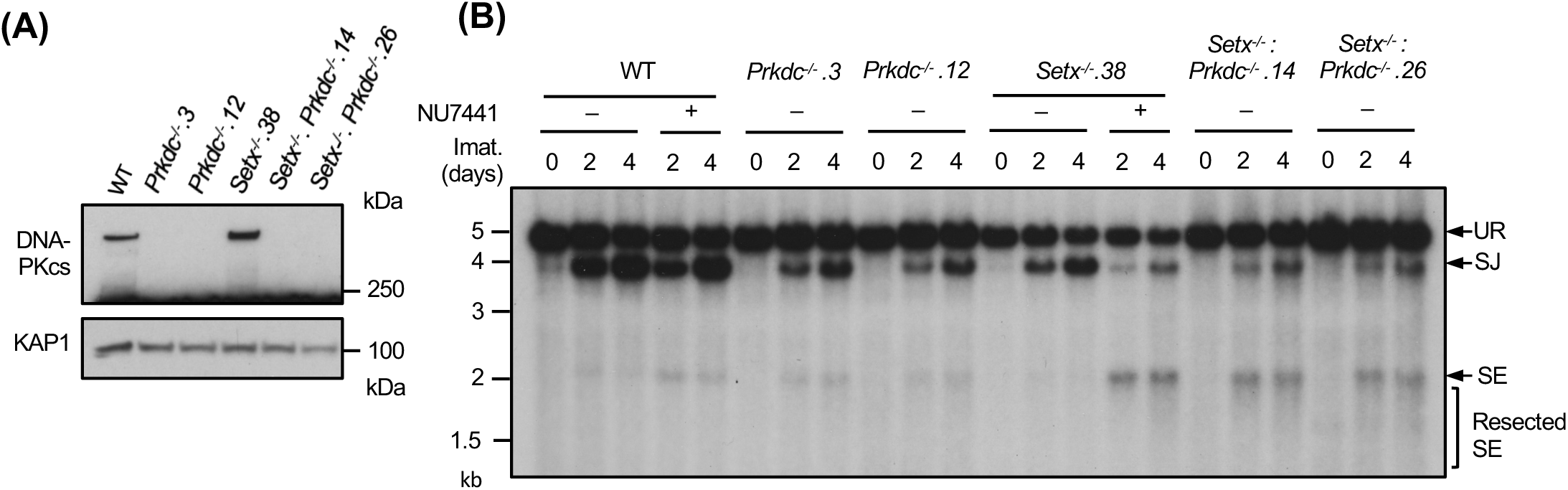
Combined loss of senataxin and DNA-PKcs proteins impairs NHEJ-mediated RAG DSB repair. (A) Western blot analysis of cell lysates from WT, *Prkdc*^-/-^, *Setx*^-/-^ and *Setx*^-/-^: *Prkdc*^-/-^ abl pre-B cells using DNA-PKcs and GAPDH antibodies. (B) Southern blot analysis of *EcoR*V-digested genomic DNA isolated from WT, *Prkdc*^-/-^, *Setx*^-/-^ and *Setx*^-/-^: *Prkdc*^-/-^ abl pre-B cells with pMX-DE^SJ^ and treated with imatinib (imat.) in the presence or absence of the DNA-PKcs inhibitor NU7441 for the indicated times. R. SE indicates resected SE restriction fragments.

### Senataxin helicase activity is required for NHEJ

Senataxin has two structured domains: the N-terminal domain and the helicase domain residing in the carboxyl terminus (*25, 26, 31*). The helicase activity of senataxin is critical for its known biological functions including transcriptional termination, gene expression and R-loop resolution (*25-27, 32, 33*). We therefore determined the impact of the senataxin helicase domain in RAG DSB repair. To circumvent the lack of reliable antibodies to murine senataxin, we inserted a triple HA epitope tag (3HA) at the 3’ end of both *Setx* alleles in frame with the *Setx* coding sequence in WT abl pre-B cells, termed *Setx^3HA/3HA^* hereafter (Figs. S3A and S3B). V(D)J recombination in *Setx^3HA/3HA^* abl pre-B cells was comparable to WT abl pre-B cells and inactivation of senataxin-3HA in *Setx^3HA/3HA^* abl pre-B cells by CRISPR/Cas9 led to V(D)J recombination defects similar to clonal *Setx*^-/-^ abl pre-B cells upon DNA-PKcs kinase inhibition (Figs. S3A, S3C and 1D).

To assess the contribution of the senataxin helicase domain to NHEJ, we introduced homozygous K1945R and L2131W mutations in senataxin that abolish the ATPase and RNA binding activities of senataxin, respectively (*25-27, 32*). The K1945R and L2132W mutations did not result in a change in the abundance of the mutant proteins when compared to WT senataxin (Figs. 4A, S4A and S4C). Upon imatinib treatment, both *Setx^K1945R/K1945R^*and *Setx^L2132W/L2132W^* abl pre-B cells formed SJs to an extent similar to WT and *Setx*^-/-^ abl pre-B cells (Figs. 4B, S4B and S4D). However, when these mutant cells were treated with imatinib and the DNA-PKcs kinase inhibitor NU7441, they exhibited a significant reduction in SJ formation and an increased accumulation of unrepaired SEs (Figs. 4B, S4B and S4D), similar to that observed in *Setx*^-/-^ abl pre-B cells (Figs. 2D and S1B). We conclude that an intact helicase function of senataxin is required for NHEJ in the absence of DNA-PKcs kinase activity.

**Figure 4:**
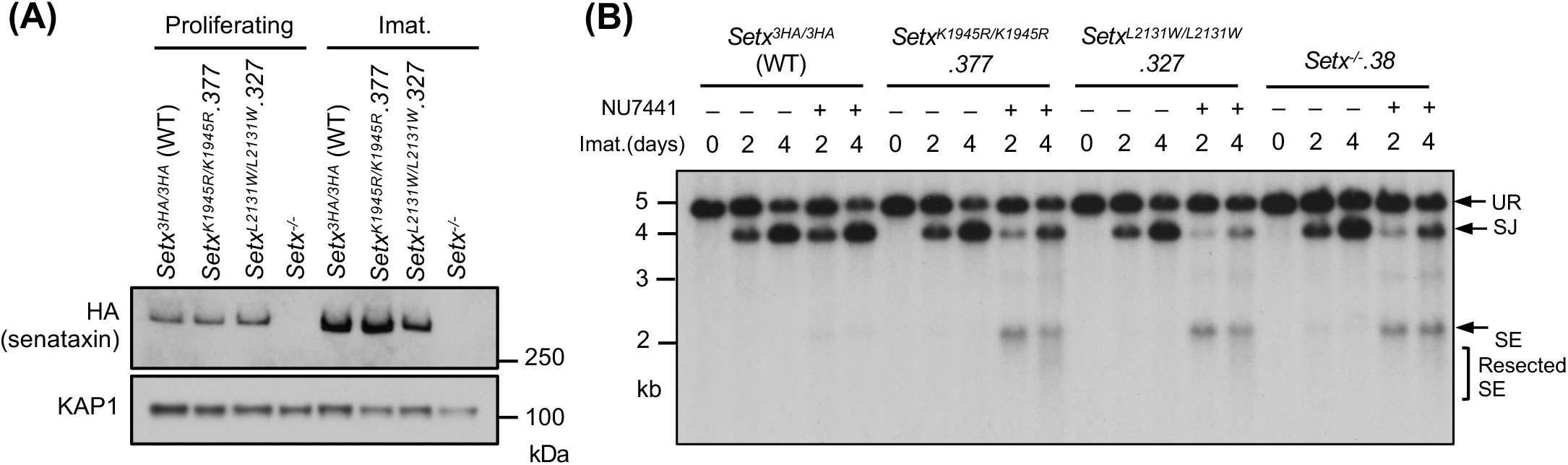
Senataxin helicase activity is required for RAG DSB repair. (A) Western blot analysis of cell lysates from proliferating or imatinib (imat.)-treated *Setx^-/-^*, *Setx^3HA/3HA^* and *Setx^K1945R/K1945R^*and *Setx^L2131W/L2131W^* abl pre-B cells using HA and KAP1 antibodies. (B) Southern blot analysis of *EcoR*V-digested genomic DNA from cells described in (*A*) with pMX-DEL^SJ^ and treated with imatinib (imat.) in the presence or absence of the DNA-PKcs kinase inhibitor NU7441 for the indicated times.

### Senataxin NHEJ function is redundant with the RECQL5 and HLTF helicases

Our results suggest that senataxin acts in the same pathway as ATM to promote NHEJ when DNA-PKcs is absent. To test whether other helicases might function in NHEJ and be similarly governed by DNA-PKcs, we conducted a genome-wide CRISPR/Cas9 gRNA screen in *Setx^-^*^/-^ abl pre-B cells containing the pMG-INV retroviral recombination substrate. This revealed that gRNAs to *Recql5*, encoding the RECQ helicase family member RECQL5, were enriched in the GFP^-^ population of *Setx^-^*^/-^, but not WT, abl pre-B cells (Tables S2 and S3). Loss of RECQL5 alone did not have a discernible impact on V(D)J recombination as imatinib-treated *Recql5*^-/-^ and WT cells had a similar percentage of GFP^+^ cells (Figs. 5A and 5B). Analyses of *Setx*^-/-^: *Recql5*^-/-^ abl pre-B cell lines, however, revealed a consistent reduction in the percentage of GFP^+^ cells as compared to *Setx*^-/-^ abl pre-B cells (Figs. 5C and 5D). The V(D)J recombination deficiency in *Setx*^-/-^: *Recql5*^-/-^ abl pre-B cells is a result of a defect in NHEJ-mediated RAG DSB repair as evidenced by Southern blot analysis of pMX-DEL^SJ^ in these cells, which revealed an accumulation of unrepaired SEs (Fig. 5E). Taken together, these data show that loss of RECQL5 in *Setx*^-/-^ abl pre-B cells compromises NHEJ.

**Figure 5:**
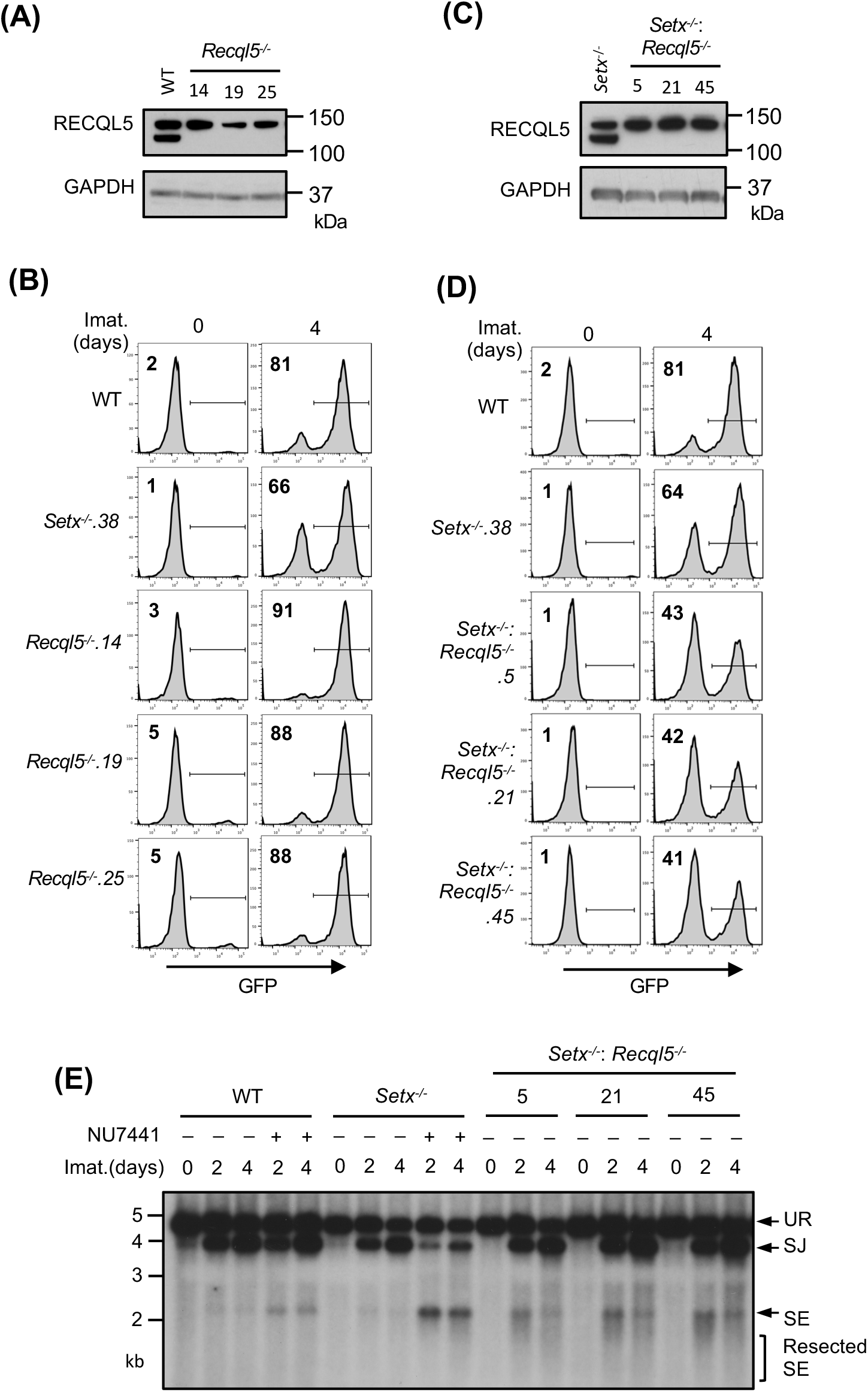
Loss of RECQL5 modestly impairs RAG-DSB repair in *Setx*^-/-^ abl pre-B cells. (A) Western blot analysis of cell lysates from WT and three independently isolated *Recql5*^-/-^ abl pre-B cells using RECQL5 and GAPDH antibodies. (B) Flow cytometric analysis for GFP expression in cells described in (A) with pMG-INV and treated with imatinib (imat.) for the indicated times. The percentages of GFP^+^ cells are indicated in the top left corners of the histograms. (C) Western blot analysis of cell lysates from *Setx*^-/-^ and three independently isolated *Setx*^-/-^: *Recql5*^-/-^ abl pre-B cells using RECQL5 and GAPDH antibodies. (D) Flow cytometric analysis for GFP expression in cells described in (C) treated with imatinib for the indicated times. (E) Southern blot analysis of *EcoR*V-digested genomic DNA from cells described in (*D*) with retroviral pMX-DEL^SJ^ and treated with imatinib (imat.) in the presence or absence of the DNA-PKcs kinase inhibitor NU7441 for the indicated times.

We reasoned that the modest impact of combined loss of senataxin and RECQL5 on NHEJ-mediated RAG repair, as compared to co-inactivation of senataxin and DNA-PKcs, could be due to the activity of additional helicases. Indeed, a genome-wide CRISPR/Cas9 gRNA screen in imatinib-treated *Setx*^-/-^*: Recql5*^-/-^ abl pre-B cells identified multiple gRNAs to *Hltf*, encoding the double strand DNA translocase and chromatin remodeler HLTF, enriched in the GFP^-^ population (Table S4). Loss of HLTF, in WT or *Setx*^-/-^ abl pre-B cells had no effect on V(D)J recombination as revealed by flow cytometric analysis of cells containing pMG-INV (Fig. S5). Moreover, abl pre-B cells lacking both RECQL5 and HLFT also exhibited normal V(D)J recombination (Fig. S6). However, *Setx*^-/-^*: Recql5*^-/-^: *Hltf*^-/-^ abl pre-B cells exhibited a substantial reduction in GFP^+^ cells upon induction of V(D)J recombination when compared to *Setx*^-/-^ and *Setx*^-/-^*: Recql5*^-/-^ abl pre-B cells (Figs. 6A, 6B and 5D). Reduced SJ formation and increasing levels of unrepaired SEs were also observed upon Southern blot analyses of pMX-DEL^SJ^ indicating a significant NHEJ defect in *Setx*^-/-^*: Recql5*^-/-^: *Hltf*^-/-^ abl pre-B cells (Fig. 6C). We conclude that the RECQL5 and HLTF helicases function redundantly with senataxin during the NHEJ-mediated repair of RAG DSBs. In this regard, to test whether RECQL5 and HLTF could exert their function as a complex, we conducted immunoprecipitation assay in imatinib-treated abl pre-B cells expressing HA-tagged RECQL5. Our result showed that in the HA antibody immunocomplex, both HA-RECQL5 and endogenous HLTF could be detected in a DNA damage-independent manner, indicating that a fraction of RECQL5 and HLTF constitutively associate with each other in cells (Fig. 6D).

**Figure 6:**
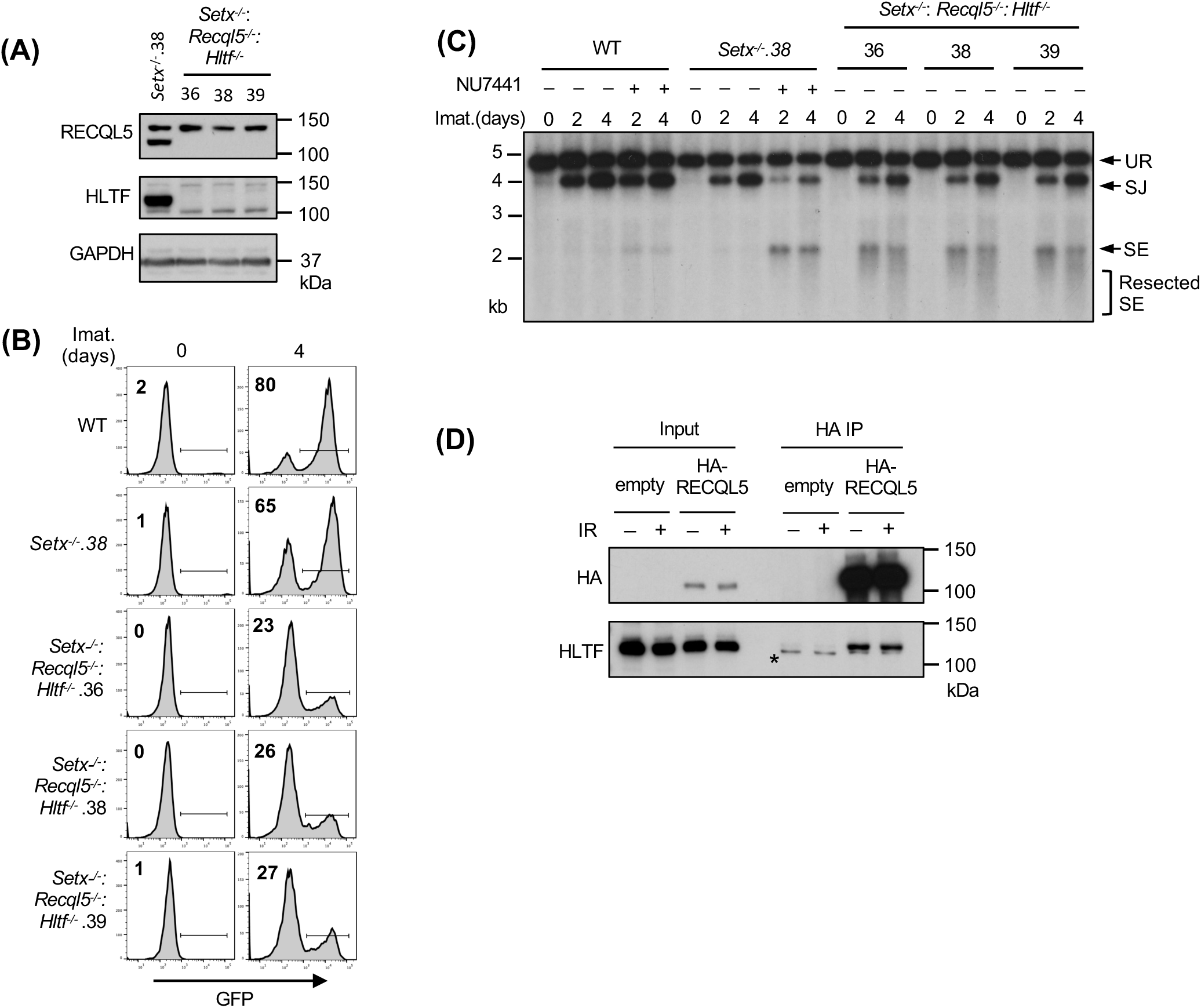
Combined loss of senataxin, RECQL5 and HLTF results in defected RAG DSB repair. (A) Western blot analysis of cell lysates from *Setx*^-/-^ and three independently isolated *Setx*^-/-^: *Recql5*^-/-^*: Hltf*^-/-^ abl pre-B cells using RECQL5, HLTF and GAPDH antibodies. (B) Flow cytometric analysis for GFP expression in WT, *Setx*^-/-^ and *Setx*^-/-^: *Recql5*^-/-^*: Hltf*^-/-^ abl pre-B cells with pMG-INV and treated with imatinib (imat.) for the indicated times. The percentages of GFP^+^ cells are indicated in the top left corners of the histograms. (C) Southern blot analysis of *EcoR*V-digested genomic DNA from abl pre-B cells described in (*B*) with pMX-DE^SJ^ and treated with imatinib (imat.) in the presence or absence of the DNA-PKcs kinase inhibitor NU7441 for the indicated times. (D) Western blot analysis of HLTF co-immunoprecipitation with ectopically expressed HA-RECQL5 in imatinib-treated abl pre-B cell lysate with HA and HLTF antibodies. The asterisk indicates the non-specific recognizing bands by the HLTF antibody.

### ATM activity is required for RAG DSB repair in *Recql5*^-/-^: *Hltf^-^*^/-^ abl pre-B cells

Our results thus far suggest that two pathways, one regulated by senataxin and the other by RECQL5 and HLTF, redundantly support NHEJ-mediated RAG DSB repair. Moreover, that inhibition of the kinase activity of DNA-PKcs, but not ATM, uniquely impairs RAG DSB repair in *Setx*^-/-^ abl pre-B cells suggests an epistatic relationship between ATM and senataxin. We next determined whether DNA-PKcs could function with RECQL5 and HLTF in a similar manner. Inhibition of DNA-PKcs kinase activity by NU7441, despite strongly blocking V(D)J recombination in *Setx*^-/-^ abl pre-B cells, only had modest effects in *Recql5^-/-^*, *Hltf^-/-^*and *Recql5*^-/-^: *Hltf*^-/-^ abl pre-B cells as measured by GFP expression from pMG-INV in a flow cytometric assay (Figs. 7A, S5B and S7). On the contrary, inhibition of ATM kinase activity with KU55933 slightly decreased V(D)J recombination in WT, *Setx*^-/-^, *Recql5* ^-/-^and *Hltf^-/-^*abl pre-B cells but substantially reduced the percentage of GFP^+^ cells in *Recql5*^-/-^: *Hltf*^-/-^ abl pre-B cells (Figs. 7A, S5B and S7). Southern blot analyses of KU55933-treated *Recql5*^-/-^: *Hltf*^-/-^ abl pre-B cells also revealed the accumulation of unrepaired SE fragments, indicating defective RAG DSB repair in these cells (Fig. 7B). Taken together, these results suggest that ATM may regulate senataxin activity and DNA-PKcs may regulate the activities of RECQL5 and HLTF to promote NHEJ-mediated DSB repair.

**Figure 7:**
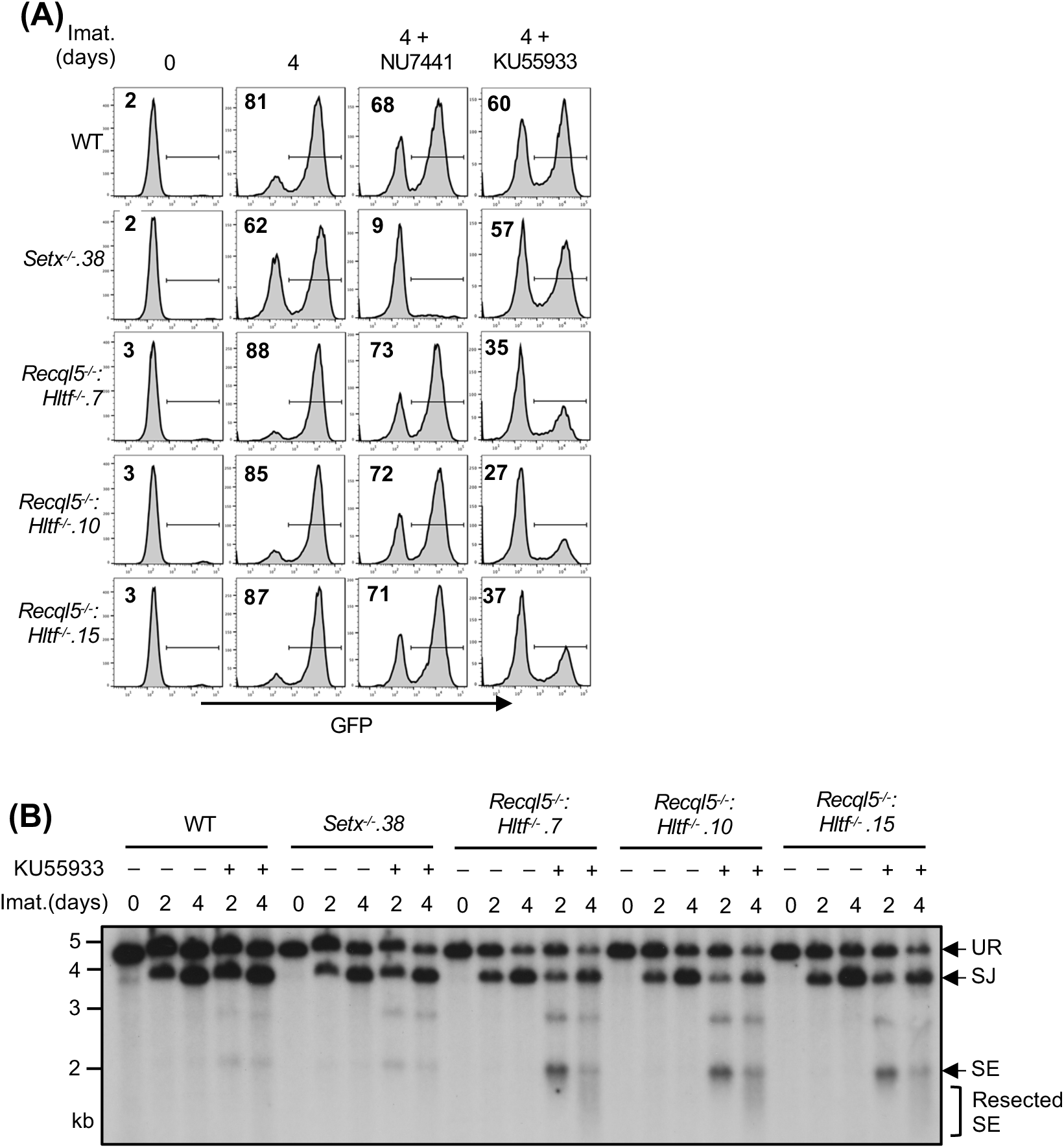
ATM activity is required for RAG DSB repair in *Recql5*^-/-^: *Hltf^-^*^/-^ abl pre-B cells. (A) Flow cytometric analysis for GFP expression from pMG-INV in WT, *Setx^-^*^/-^ and three independently isolated *Recql5*^-/-^: *Hltf^-^*^/-^ abl pre-B cell lines treated with imatinib (imat.) in the presence or absence of the DNA-PKcs kinase inhibitor NU7441 or the ATM kinase inhibitor KU55933 for the indicated times. The percentages of GFP^+^ cells are indicated in the top left corners of the histograms. (B) Southern blot analysis of *EcoR*V-digested genomic DNA from cells described in (*A*) with pMX-DE^SJ^ and treated with imatinib (imat.) in the presence or absence of KU55933 for the indicated times.

### Partial inhibition of RNA polymerase II activity improves RAG DSB repair in helicase-deficient abl pre-B cells

During V(D)J recombination, RAG DSBs are generated and repaired in actively transcribed chromatin regions. Given mounting evidence demonstrating the impact of transcription on DSB repair, we tested whether the NHEJ defect seen in our mutant abl pre-B cells is transcription-dependent (*34, 35*). To this end, *Setx^-^*^/-^ abl pre-B cells were exposed to low doses of THZ1, a highly selective inhibitor to the RNA pol II CTD kinase CDK7, 24 hours after imatinib and DNA-PKcs inhibitor treatment (*36*). In Southern blot analyses, we observed that with increasing concentrations of THZ1, the intensity of the restriction fragments corresponding to SEs decreased. Indeed, quantifications showed that the percentage of unrepaired SEs in the total amount of the recombination substrate cleaved by RAG (the sum of intensities of restriction fragments to repaired SJs and unrepaired SEs) was reduced in cells treated with increasing concentrations of THZ1 (Figs. 8A and S8A). Similar observations were made in *Setx*^-/-^*: Recql5*^-/-^: *Hltf*^-/-^ and ATM kinase inhibitor-treated *Recql5*^-/-^: *Hltf*^-/-^ abl pre-B cells (Figs. 8B, 8C, S8B and S8C). Therefore, these results indicate that partial inhibition of RNA pol II activity in abl pre-B cells can partly rescue NHEJ defects due to combined deficiency of senataxin and DNA-PKcs, co-inactivation of ATM, RECQL5 and HLTF, and loss of senataxin, RECQL5 and HLTF.

**Figure 8:**
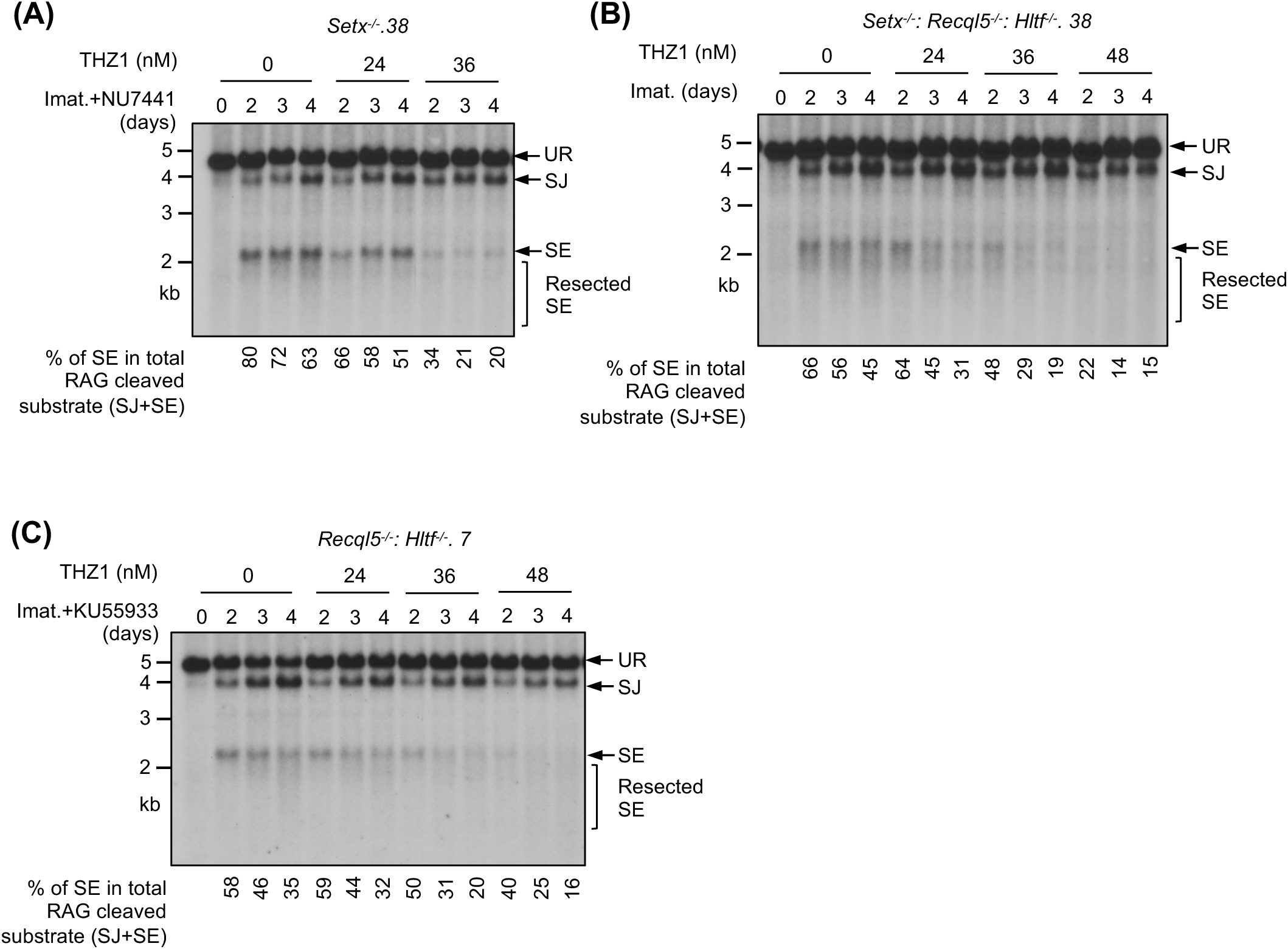
Partial inhibition of RNA polymerase II improves NHEJ-mediated RAG DSB repair in helicase-deficient abl pre-B cells. Southern blot analysis of *EcoR*V-digested genomic DNA from pMX-DEL^SJ^ containing, imatinib (imat.) treated *Setx*^-/-^ (with the DNA-PKcs kinase inhibitor NU7441) (A), *Setx*^-/-^: *Recql5*^-/-^*: Hltf*^-/-^ (B) and *Recql5*^-/-^*: Hltf*^-/-^ (with the ATM kinase inhibitor KU55933) (C) abl pre-B cells with increasing concentrations of the CDK7 inhibitor THZ1 for the indicated times. The percentages of unrepaired (full length and resected) SEs in total RAG cleaved recombination substrates (SE+SJ) are shown below the Southern blot images.

### Aberrant DNA end joining in abl pre-B cells deficient in senataxin, RECQL5 and HLTF

In *Setx*-deficient abl pre-B cells treated with DNA-PKcs inhibitor, not only did we observe the accumulation of unrepaired SEs and CEs in Southern blot analyses, the restriction fragments of these unrepaired DNA ends were heterogeneous in size, suggesting that they were aberrantly resected (Figs. 2, 4, S1 and S4) (*37-40*). Similar observations were made in *Setx*^-/-^*: Recql5*^-/-^, *Setx*^-/-^*: Recql5*^-/-^: *Hltf*^-/-^ and ATM kinase inhibitor-treated *Recql5*^-/-^: *Hltf*^-/-^ abl pre-B cells (Figs. 5E, 6C, 7B). Indeed, when MRE11, which initiates DNA resection, was depleted by CRISPR/Cas9, the heterogeneity in size of SE bands was markedly reduced and a single strong band appeared, consistent with the notion that the unrepaired DNA ends are resected in these cells (Figs. 9A and 9B) (*41*).

**Figure 9:**
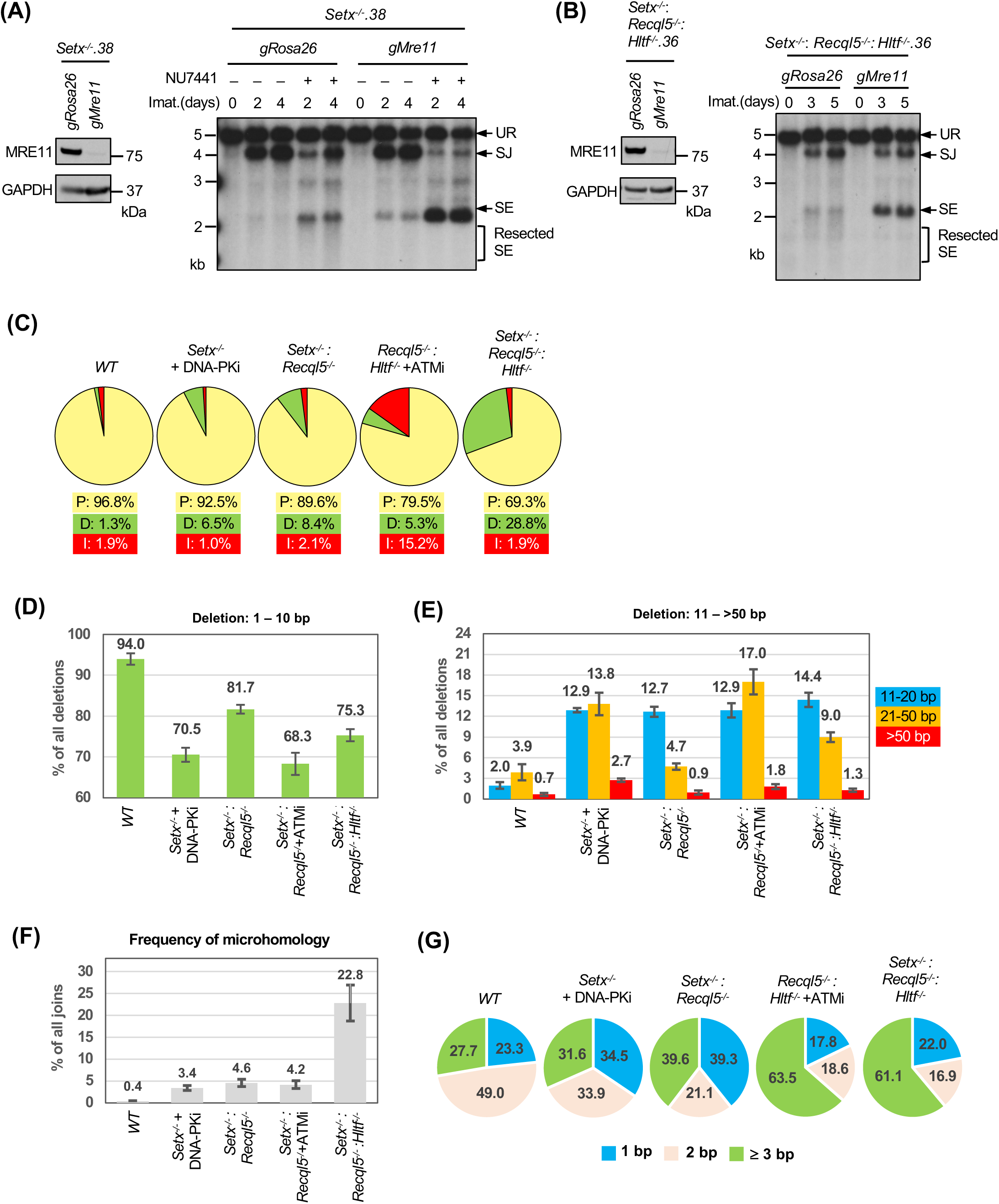
Aberrant DNA end joining in abl pre-B cells deficient in senataxin, RECQL5 and HLTF. (A, B) Left, western blot analysis of cell lysate from *Setx*^-/-^ *(A)* or *Setx*^-/-^: *Recql5*^-/-^*: Hltf*^-/-^ *(B)* abl pre-B cells expressing Cas9 and gRNAs to *Rosa26* (*gRosa26*) or *Mre11* (*gMre11*) using MRE11 and GAPDH antibodies. Right, Southern blot analysis of *EcoR*V-digested genomic DNA from the aforementioned cells with pMX-DEL^SJ^ and treated with imatinib (imat.) in the presence or absence of the DNA-PKcs kinase inhibitor NU7441 for the indicated times. (C) The percentages of precise joins (P, yellow), joins with deletions (D, green) and joins with insertions (I, red) in SJs amplified from pMG-INV in imatinib treated WT, *Setx*^-/-^ (with NU7441), *Setx*^-/-^*: Recql5*^-/-^, *Recql5*^-/-^*: Hltf^-^*^/-^ (with the ATM kinase inhibitor KU55933) and *Setx*^-/-^*: Recql5*^-/-^*: Hltf^-^*^/-^ abl pre-B cells. (D, E) The percentages of short (1-10 bp) (D) and long (11->50 bp) (E) deletions in deletion-containing SJs amplified from pMG-INV in the indicated abl pre-B cell lines. The numbers above each bar indicate the average percentages of SJ sequences with the specific length distributions of deletion. (F) The percentages of SJs with microhomology at joining junctions in the indicated abl pre-B cell lines with the average percentages of SJs with microhomology shown on top of each bar. (G) The microhomology length distribution (% of all microhomology) of the SJs shown in (*F*). Number of samples sequenced per genotype: WT: 7, WT + NU7441: 3, *Setx^-/-^*: 6, *Setx*^-/-^ + NU7441: 3, *Recql5^-/-^*: 3, *Hltf*^-/-^: 3, *Setx*^-/-^: *Recql5*^-/-^: 4, *Setx*^-/-^: *Hltf*^-/-^: 3, *Recql5*^-/-^: *Hltf^-/-^*: 3, *Recql5*^-/-^: *Hltf^-^*^/-^ + KU55933: 3, *Setx*^-/-^: *Recql5*^-/-^: *Hltf*^-/-^: 4.

During NHEJ-mediated DSB repair, the resection of DNA ends and the potential generation of single stranded overhangs could lead to aberrant homology-mediated joining, which frequently results in nucleotide loss. During V(D)J recombination, accurate SJ formation occurs with only a small fraction of SJs losing one or two nucleotides (*42*). To survey the structural variants of SJs in cells deficient in ATM, DNA-PKcs, senataxin, RECQL5 and HLTF, we PCR amplified and sequenced pMG-INV SJs using genomic DNA extracted from different abl pre-B cell lines. As expected, the majority of the SJs in WT abl pre-B cells were formed precisely (∼97%) with a small percentage of nucleotide loss (deletion) and gain (insertion) (Fig. 9C and Table S5). Loss of senataxin or inhibition of DNA-PKcs by NU7441 individually led to a small increase in the fraction of SJs containing deletions (Table S5). However, the inhibition of DNA-PKcs in *Setx^-/-^* abl pre-B cells resulted in a pronounced increase in the frequency of SJs containing deletions (Fig. 9C and Table S5). Higher levels of SJs with significant deletions were also observed in *Setx^-/-^: Recql5*^-/-^, ATM inhibitor-treated *Recql5*^-/-^: *Hltf^-^*^/-^ and *Setx*^-/-^*: Recql5*^-/-^: *Hltf*^-/-^ abl pre-B cells (Fig. 9C and Table S5). Most of the deletions are shorter than 10 bp; however, longer deletions (>10 bp) are more frequent in the mutants in which higher fractions of SJs contain deletions (Figs. 9D and 9E). Notably, in addition to deletions, a significant fraction of SJs in ATM-inhibited *Recql5*^-/-^: *Hltf^-^*^/-^ abl pre-B cells also contained insertions (Fig. 9C and Table S5).

Sequence analyses revealed that less than 1% of SJs in WT abl pre-B cells exhibit microhomology at SJs whereas 23% of SJs from *Setx*^-/-^*: Recql5*^-/-^: *Hltf*^-/-^ abl pre-B cells exhibit microhomology with 60% of these exhibiting 3 base pairs or more of microhomology (Fig. 9F and 9G). We conclude that combined deficiency of DNA-PKcs and senataxin, co-inactivation of ATM, RECQL5 and HLTF and loss of senataxin, RCQL5 and HLTF result in an increase in aberrant joining during NHEJ and that this joining is more dependent on the use of microhomologies.

## Discussion

In this study, we exploited V(D)J recombination, a process requiring NHEJ for proper completion, to investigate novel factors that function in ATM- or DNA-PKcs-pathways to promote NHEJ. Our findings show that a group of distinct helicases and a dsDNA translocase, senataxin, RECQL5 and HLTF, redundantly facilitate NHEJ-mediated RAG DSB repair during V(D)J recombination with senataxin likely functioning in an ATM-dependent manner while RECQL5 and HLTF functioning in a DNA-PKcs dependent pathway. Senataxin and RECQL5 have been implicated in DSB repair while the known functions of HLTF are largely restricted to the response to replicative stress (*43-50*). Senataxin, RECQL5 and HLTF are all able to resolve unique DNA structures *in vivo* or *in vitro*. Senataxin can unwind RNA:DNA hybrids, including R-loops, and loss of senataxin can lead to R-loop accumulation at DSBs (*44*). The helicase activity of RECQL5 is important for resolving Holiday junction-like structures *in vitro* and HLTF has recently been shown to promote the resolution of noncanonical nucleic acid structures known as G-quadruplexes (G4s) (*51, 52*). Given the activities of these helicases in untangling unique nucleic acid structures, it is possible that they may function redundantly to ensure that inhibitory structures do not accumulate at DSB ends to impede the recruitment of NHEJ factors or prevent DNA end ligation (*46*). Notably, RECQL5 and HLTF can also remodel or remove chromatin-bound proteins to facilitate proper duplication of common fragile sites in mitosis and replication fork reversal, respectively (*53, 54*). Moreover, recent studies demonstrated that HLTF disrupts the Cas9-DNA postcleavage complex prior to DSB repair (*55, 56*). Therefore, in addition to inhibitory DNA structures, it is possible that at least some of these proteins could function for timely removal of proteins bound to DSB ends. Along these lines, RNA pol II-dependent transcription is often accompanied by non-canonical nucleic acid structures such as R-loops and G4s and aberrant accumulation of these structures leads to DNA damage (*57, 58*). Our finding that RAG DSB repair is more efficient when RNA pol II activity is suppressed raises the possibility of a transcription-related conflict with NHEJ that is normally prevented by concerted activities of senataxin, RECQL5 and HLTF.

ATM and DNA-PKcs are known to phosphorylate many unique and overlapping protein targets with diverse functions that promote the DNA damage response (*13, 59*). This could in part explain the observation that ATM and DNA-PKcs promote NHEJ through pathways with overlapping functions such that cells retain considerable DNA DSB repair capability when one of these kinases is inhibited (*18-20*). Indeed, our results indicate that ATM and DNA-PKcs support RAG DSB repair through proteins with distinct helicase or translocase activities in a manner in which ATM appears epistatic to senataxin and DNA-PKcs appears epistatic to RECQL5 and HLTF. This is supported by co-inactivation of ATM, RECQL5 and HLTF, or combined deficiency in DNA-PKcs and senataxin, lead to defective RAG DSB repair. Additionally, inactivation of DNA-PKcs and ATM in *Recql5*^-/-^: *Hltf*^-/-^ and *Setx*^-/-^ abl pre-B cells, respectively, did not result in additional RAG DSB repair defects (Figs. 1, 6 and S2). We noted that DNA damage repair defects in the setting of combined deficiency in DNA-PKcs and senataxin have also been observed in fibroblasts derived from spinal muscular atrophy (SMA) patients and neurons from SMA mice, where the expression of senataxin and DNA-PKcs is compromised and elevated levels of endogenous DNA damage marked by γH2AX persist (*60*), raising the possibility that the kinase-helicase regulating pathways described here likely function in genome maintenance in a broader context. The molecular mechanisms by which ATM and DNA-PKcs regulate senataxin, RECLQ5 and HLTF during RAG DSB repair remain to be determined. It is to be noted that DNA-PKcs was reported to associate with senataxin in a previous study that also showed that senataxin localization to DSBs was diminished upon inhibition of the kinase activities of ATM and DNA-PKcs in proliferating HeLa cells (*43*). Additionally, senataxin was recently found to undergo ataxia telangiectasia and Rad3 (ATR)-dependent phosphorylation and this phosphorylation was reported to be important for senataxin association with sex chromosomes during meiosis (*61*). Therefore, senataxin can be regulated by multiple DDR kinases to execute various biological functions in a cell type-dependent manner to maintain genome integrity. It remains to be investigated whether senataxin is directly modified by these kinases and if phosphorylation fine tunes senataxin activity. Little is known regarding whether or how DDR kinases regulate the function of RECQL5 and HLTF in response to DNA damage or replicative stress. Our results provide supporting evidence for this possibility to warrant further investigation.

One of the major factors that dictate the choice of DSB repair pathways between NHEJ and HR is the structure of the broken DNA ends. Extensive resection of DNA ends generates long single strand overhangs that promote HR and limits NHEJ (*62*). To ensure that DNA DSBs to be repaired by NHEJ are not aberrantly processed, multiple DNA end protection mechanisms exist to counter the nucleolytic activity (*63, 64*). We observed that the unrepaired SEs in *Setx*^-/-^*: Recql5*^-/-^: *Hltf*^-/-^ abl pre-B cells were resected (Figs. 5C and 8B). Similar observations were also made in *Setx*^-/-^*: Recql5*^-/-^ abl pre-B cells, suggesting that at least senataxin and/or RECQL5 may suppress aberrant resection of DSBs before their joining by NHEJ. Indeed, senataxin deficiency in yeast and human cells has been reported to promote resection at DSBs (*45, 65*). Additionally, senataxin was found to interact with MRE11 in human cells, although it is not clear whether such interaction influences MRE11 nuclease activity at DSBs (*43*). In a large-scale proteomics study, senataxin was identified as a potential interactor of the pro-resection BRCA1 and the DNA end protector 53BP1 (*66*). While senataxin likely associates with BRCA1 to facilitate transcription termination and recovery from transcription-associated DNA damage, the biological implication of senataxin-53BP1 interaction remains unexplored (*67, 68*). Interestingly, MRE11 can also associate with RECQL5 in human cells and mediates the recruitment of RECQL5 to DSBs. Moreover, it was shown in an *in vitro* assay, RECQL5 could inhibit the 3’ to 5’ exonuclease activity of MRE11 (*48*). Whether RECQL5 also negatively regulates MRE11 exonuclease activity in cells are yet to be investigated. No existing evidence suggests that HLTF may directly or indirectly influence nucleases or DNA end protection proteins. We found in the survey of SJ structural variants that the loss of HLTF in *Setx*^-/-^*: Recql5*^-/-^ abl pre-B cells significantly elevated the frequency of SJs with deletions, which most likely come from nuclease-processed SEs. The lengths of deletions are also longer in *Setx*^-/-^*: Recql5*^-/-^: *Hltf*^-/-^ compared to *Setx*^-/-^*: Recql5*^-/-^: abl pre-B cells (Figs. 8C and 8D). These results suggest that HLTF may function to limit nucleolytic processing of DSBs, especially in conditions where DNA ends are prone to be resected (i.e. cells lacking senataxin and RECQL5).

Taken together our study demonstrates that a group of helicases and a translocase function redundantly to promote proper completion of NHEJ-dependent RAG DSB repair in an ATM- or DNA-PKcs-dependent manner. It is conceivable that this collection of helicases and kinases will also be important for NHEJ in other contexts such as DSB repair in G_1_/G_0_ phases of the cell cycle or the repair of DSBs in transcriptionally active regions including those generated during class switch recombination.

## Acknowledgement

Funding and core facility support:

B.P.S. is supported by National Institutes of Health grant R01 AI047829. UAB Comprehensive Flow Cytometry Core (cell sorting and flow cytometry analysis) is supported by National Institutes of Health grants P30 AI27667 and P30 CA013148 and UAB Heflin Center for Genomic Sciences (next generation sequencing) is supported by National Institutes of Health grant P30 CA013148.

## Author Contributions

B.C., A. N., J.K.T and B.P.S. designed the study and provided overall experimental guidance. B.C. and T.P. conducted the majority of the experiments and performed data analysis. Y. Z. performed sequencing data analysis. L.D.R. generated abl pre-B cell lines, performed one genome-scale CRISPR/Cas9 screens, conducted some experiments and assisted in data analysis. N.D carried out one genome-scale CRISPR/Cas9 screen. K.M., G. M-R. and J. N. assisted in some experiments.

## Data and materials availability

All data needed to evaluate the conclusions in the paper are present in the paper and/or the Supplementary Materials. Additional data or materials related to this paper may be requested from the authors.

## Supplemental Materials

### Supplemental Tables

**Table S1: CRISPR/Cas9 gRNA screen results for genes required for V(D)J recombination in WT abl pre-B cells treated with the DNA-PKcs inhibitor NU7441.**

**Table S2: CRISPR/Cas9 gRNA screen results for genes required for V(D)J recombination in *Setx*^-/-^.38 abl pre-B cells.**

**Table S3: CRISPR/Cas9 gRNA screen results for genes required for V(D)J recombination in *B.Setx*^-/-^.10 abl pre-B cells.**

**Table S4: CRISPR/Cas9 gRNA screen results for genes required for V(D)J recombination in *Setx*^-/-^: *Recql5^-/-^*.5 abl pre-B cells.**

**Table S5: Sequence analysis of SJs amplified from pMG-INV from indicated abl pre-B cells.**

**Table S6: Oligo nucleotides sequences.**

**Supplemental file 1: The reference sequence for SJ structural variant sequencing**

### Supplemental Figures

**Figure S1:**
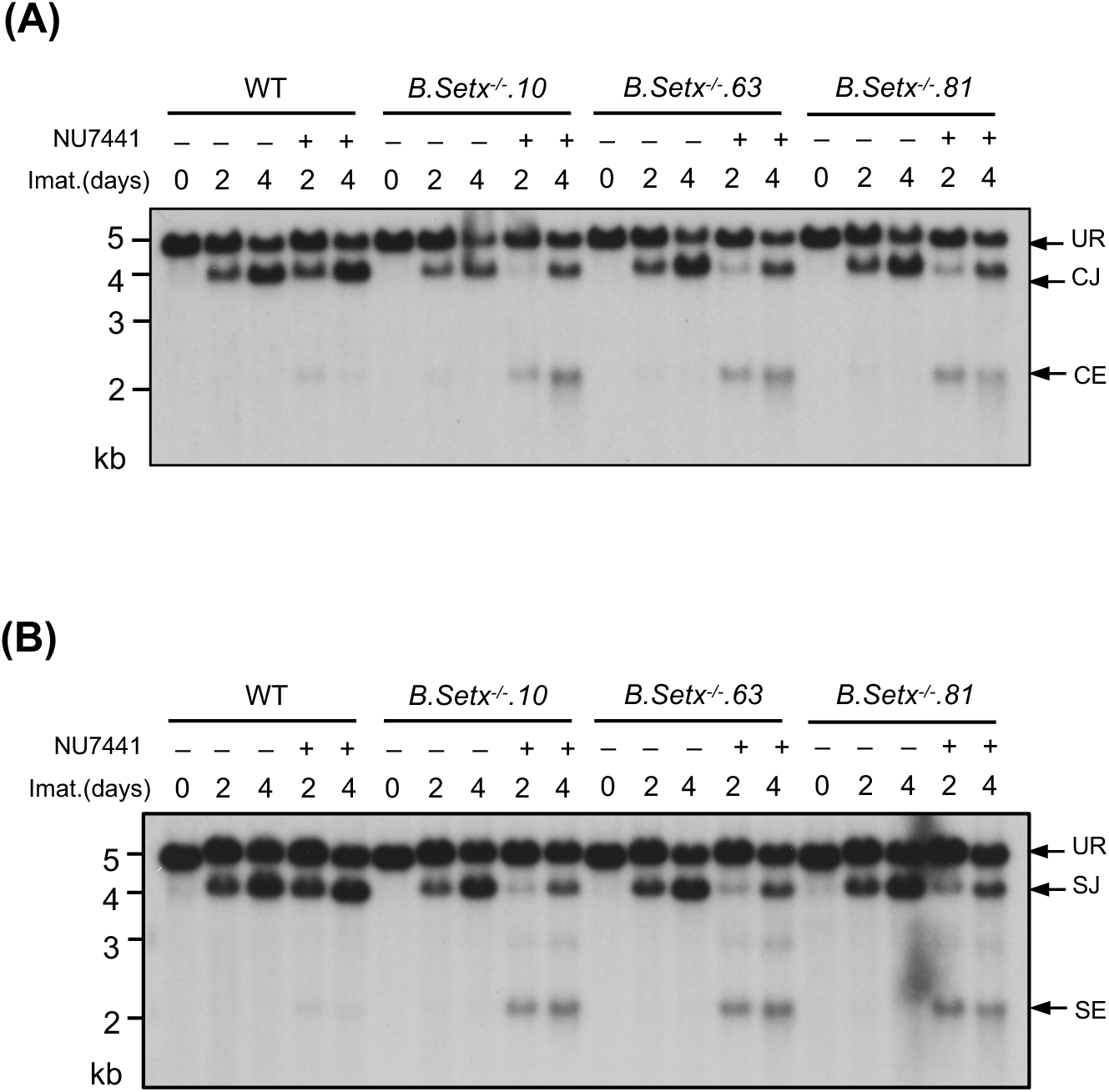
Loss of senataxin severely impairs NHEJ-mediated RAG DSB repair in DNA-PKcs-inhibited abl pre-B cells during V(D)J recombination. Southern blot analysis of *EcoR*V-digested genomic DNA from additional WT and clonal *Setx*^-/-^ abl pre-B cell lines with pMX-DEL^CJ^ (A) or pMX-DEL^SJ^ (B) treated with imatinib (imat.) in the presence or absence of the DNA-PKcs kinase inhibitor NU7441 for the indicated times.

**Figure S2:**
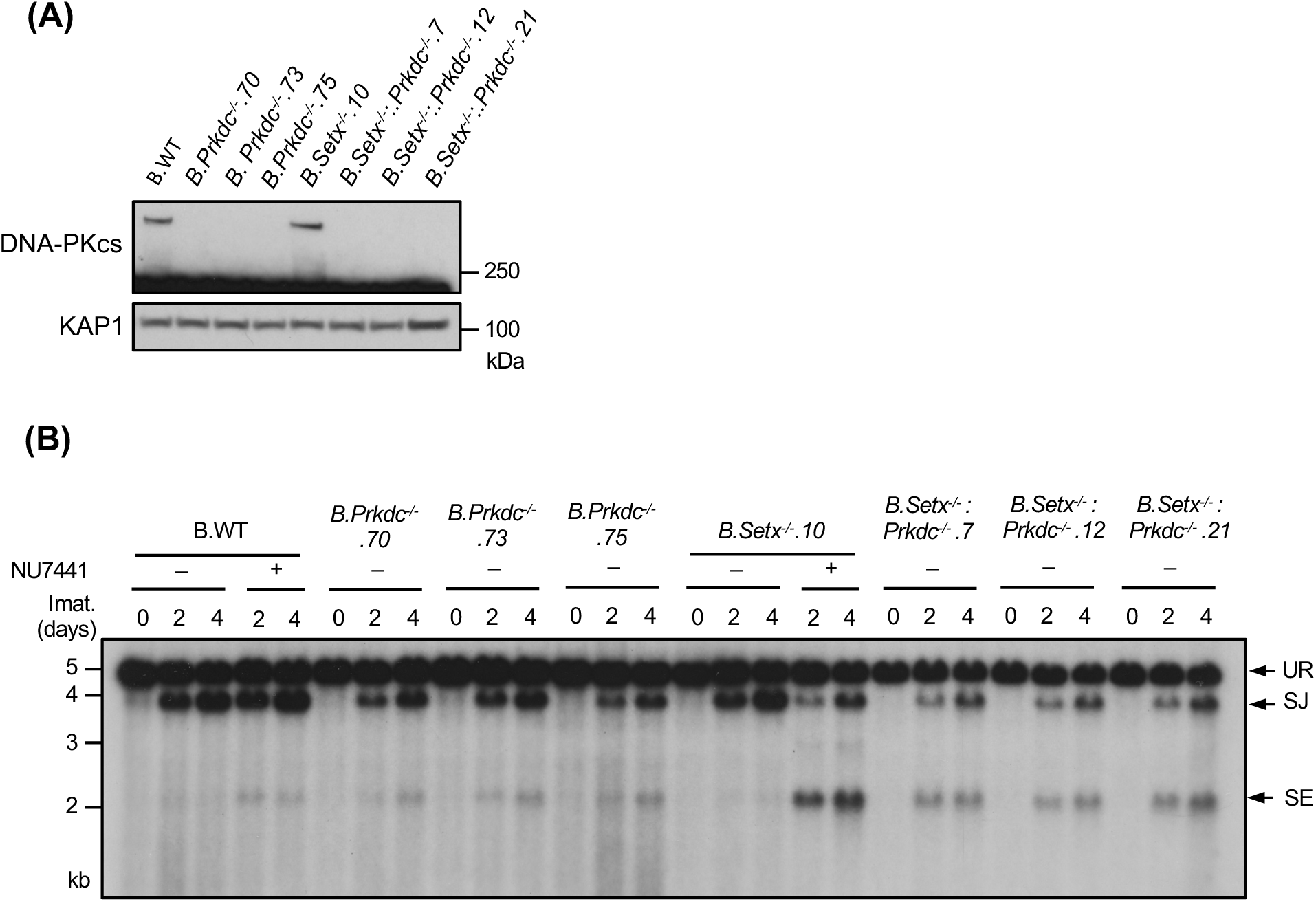
Combined loss of senataxin and DNA-PKcs proteins impairs NHEJ-mediated RAG DSB repair. (A) Western blot of cell lysates from additional WT, *Prkdc*^-/-^, *Setx*^-/-^ and *Setx*^-/-^: *Prkdc*^-/-^ abl pre-B cell lines using DNA-PKcs and GAPDH antibodies. (B) Southern blot analysis of *EcoR*V-digested genomic DNA isolated from WT, *Prkdc*^-/-^, *Setx*^-/-^ and *Setx*^-/-^: *Prkdc*^-/-^ abl pre-B cells pMX-DE^SJ^ and treated with imatinib (imat.) in the presence or absence of the DNA-PKcs inhibitor NU7441 for the indicated times.

**Figure S3:**
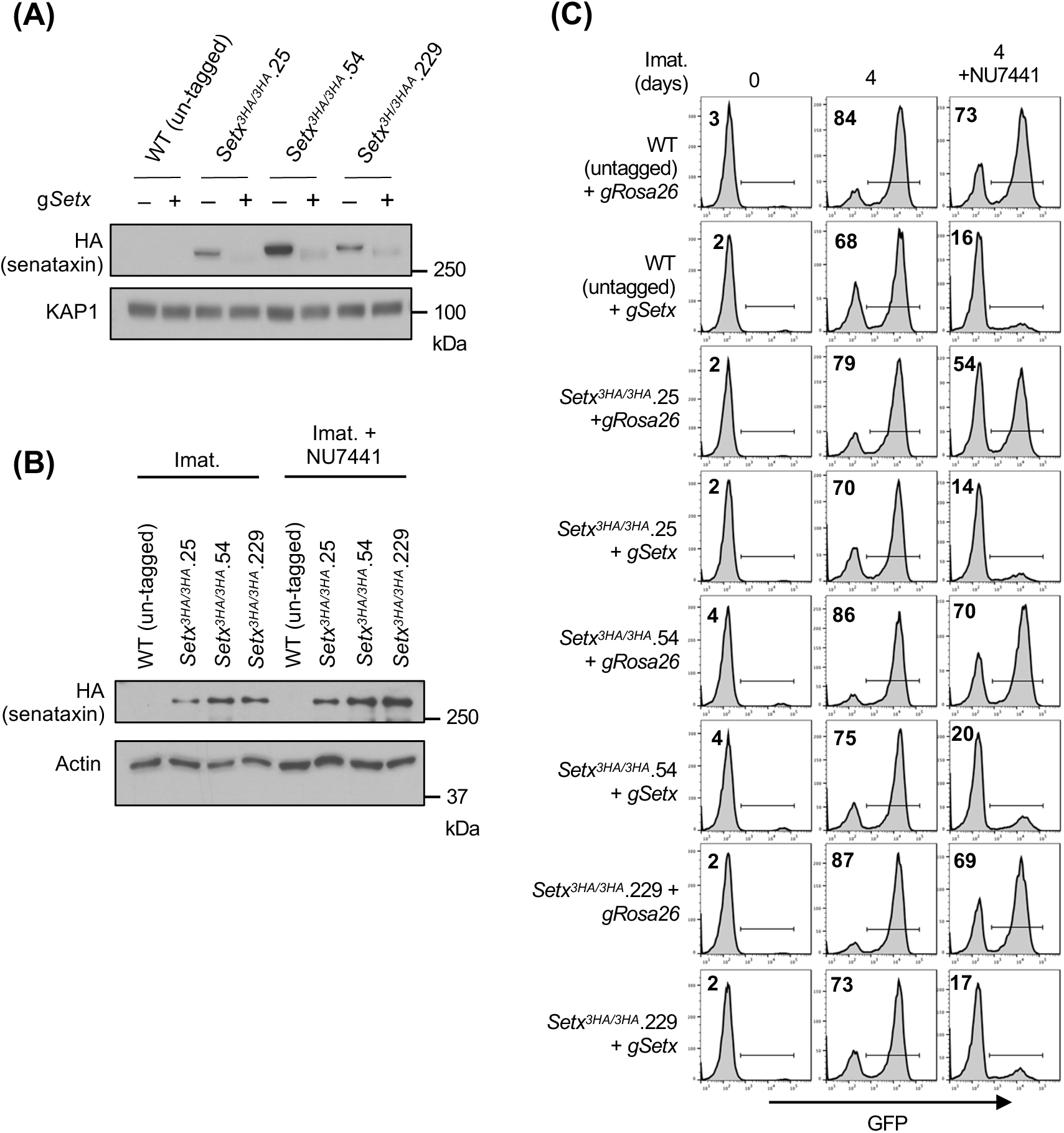
Characterization of abl pre-B cells expressing endogenous senataxin with a C-terminally tagged triple HA epitope (3HA). (A) Western blot analysis of cell lysates from proliferating WT and *Setx^3HA/3HA^* abl pre-B cells expressing Cas9 with or without Setx gRNAs (*gSetx*) using HA and KAP1 antibodies. (B) Western blot analysis of cell lysates from imatinib (imat.)-treated WT and *Setx^3HA/3HA^*abl pre-B cells with or without the DNA-PKcs kinase inhibitor NU7441 for 2 days using HA and actin antibodies. (C) Flow cytometric analysis for GFP expression from pMG-INV in WT and *Setx^3HA/3HA^* abl pre-B cells expressing Cas9 and *gRosa26* or *gSetx*, treated with imatinib in the presence or absence of NU7441 for the indicated times. The percentages of GFP^+^ cells are indicated in the top left corners of the histograms.

**Figure S4:**
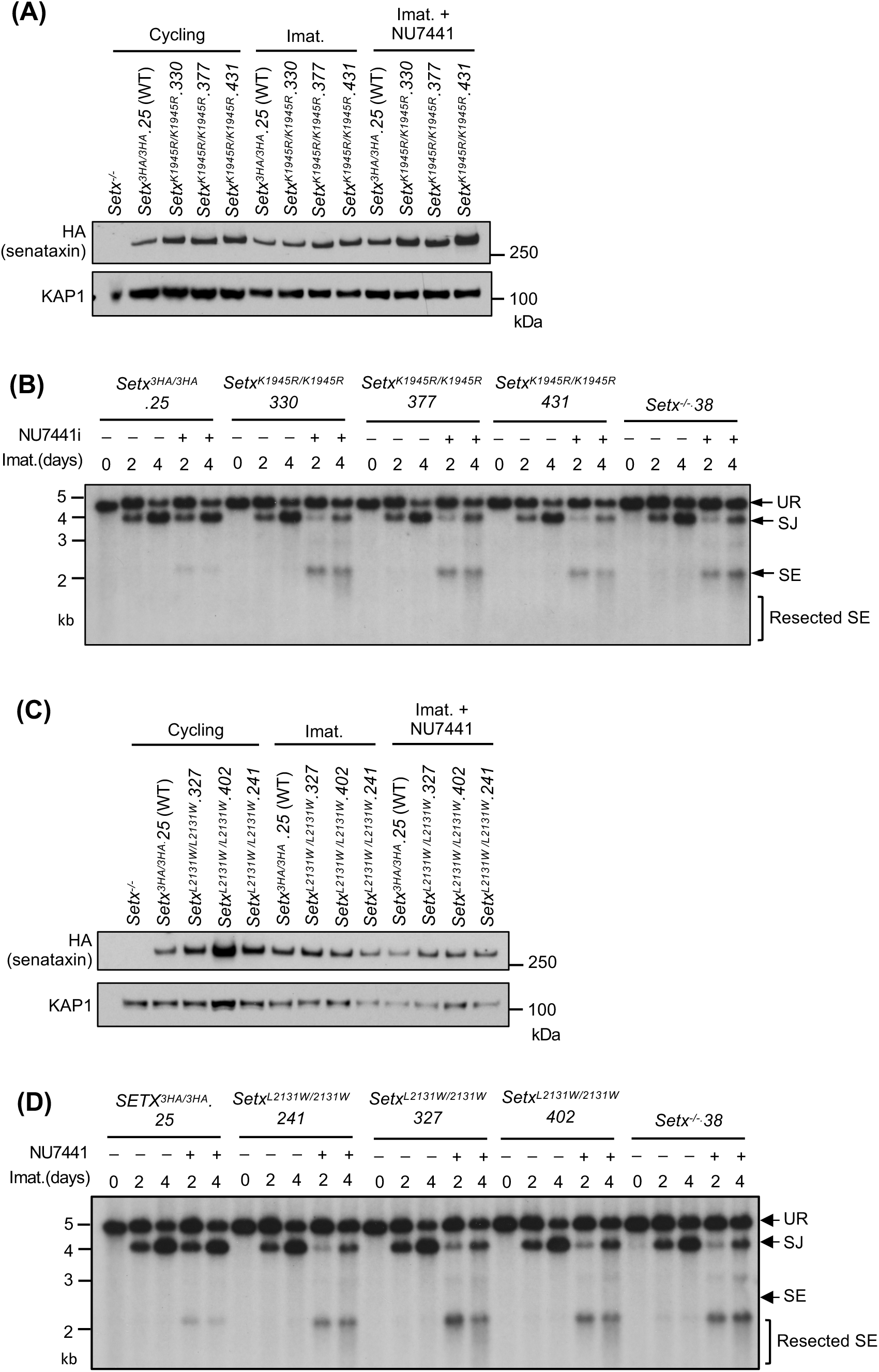
The K1945R and L2131W mutations of senataxin impair its NHEJ function. (A, C) Western blot analysis of cell lysates collected from *Setx^-^*^/-^, *Setx^3HA/3HA^*and three clones of *Setx^K1945R/K1945R^* (*A*) or *Setx^L2131W/L2131W^*(*C*) abl pre-B cells in proliferation or after treatment with imatinib (imat.) in the presence of absence of DNA-PKcs kinase inhibitor NU7441 for 2 days using HA and KAP1 antibodies. (B, D) Southern blot analysis of *EcoR*V-digested genomic DNA from *Setx^K1945R/K1945R^* (*B*) or *Setx^L2131W/L2131W^* (*D*) abl pre-B cells with pMX-DEL^SJ^ after imatinib treatment with or without NU7441 for the indicated times.

**Figure S5:**
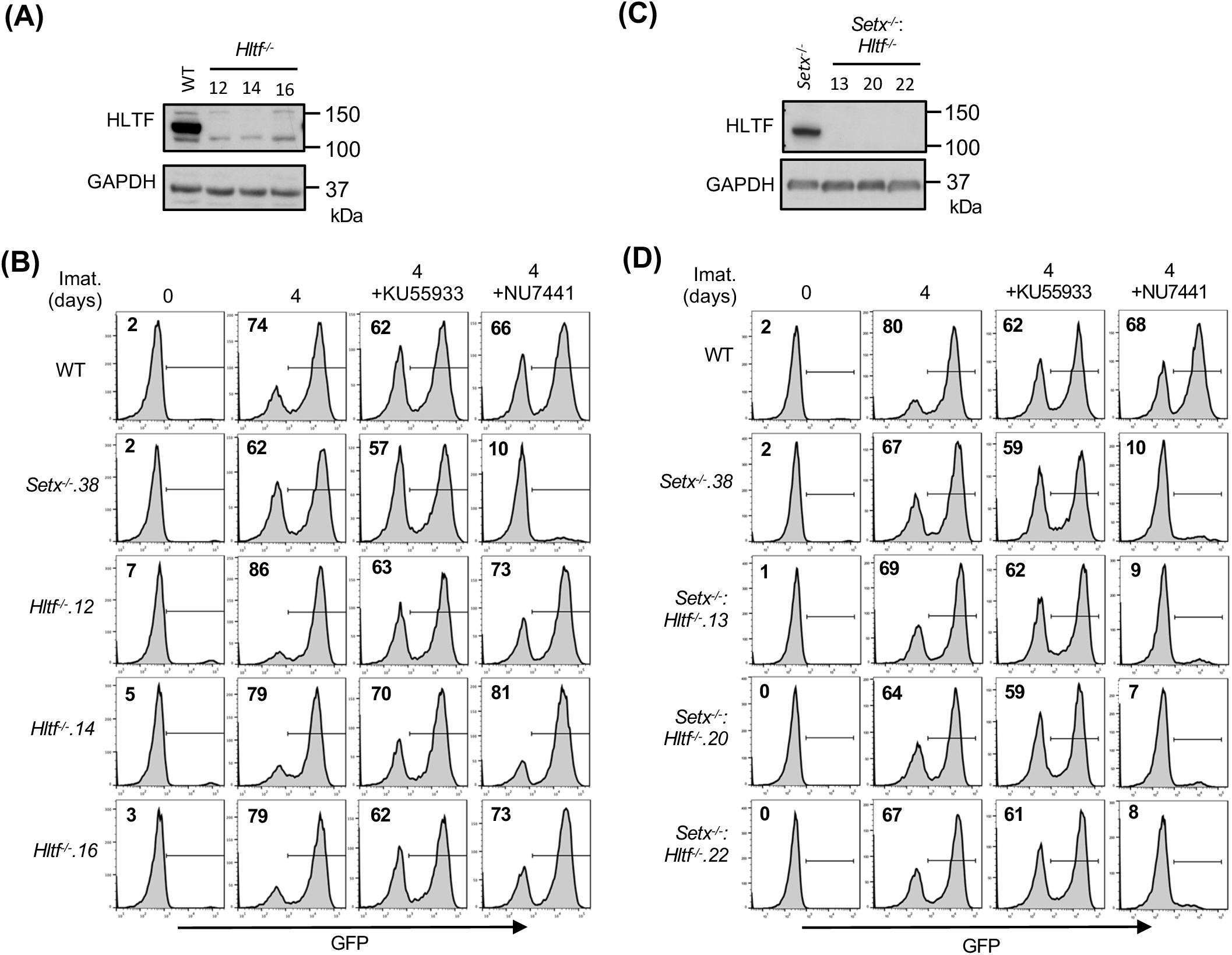
***Hltf*^-/-^ and *Setx*^-/-^: *Hltf*^-/-^ abl pre-B cells exhibit no discernible defect in V(D)J recombination.** (A) Western blot analysis of cell lysates from WT and three independently isolated *Hltf*^-/-^ abl pre-B cells using HLTF and GAPDH antibodies. (B) Flow cytometric analysis of GFP expression from pMG-INV in WT, *Setx*^-/-^ and three clones of *Hltf^-/-^* abl pre-B cells treated with imatinib alone or in the presence of the ATM kinase inhibitor KU55933 of DNA-PKcs kinase inhibitor NU7441 for the indicated times. The percentages of GFP^+^ cells are indicated in the top left corners of the histograms. (C) Western blot analysis of cell lysates from *Setx^-/-^* and three independently isolated *Setx*^-/-^: *Hltf*^-/-^ abl pre-B cells using HLTF and GAPDH antibodies. (D) Flow cytometric analysis of GFP expression from pMG-INV in WT, *Setx*^-/-^ and three clones of *Setx^-/-^*: *Hltf^-/-^* abl pre-B cells as indicated in *(B)*.

**Figure S6:**
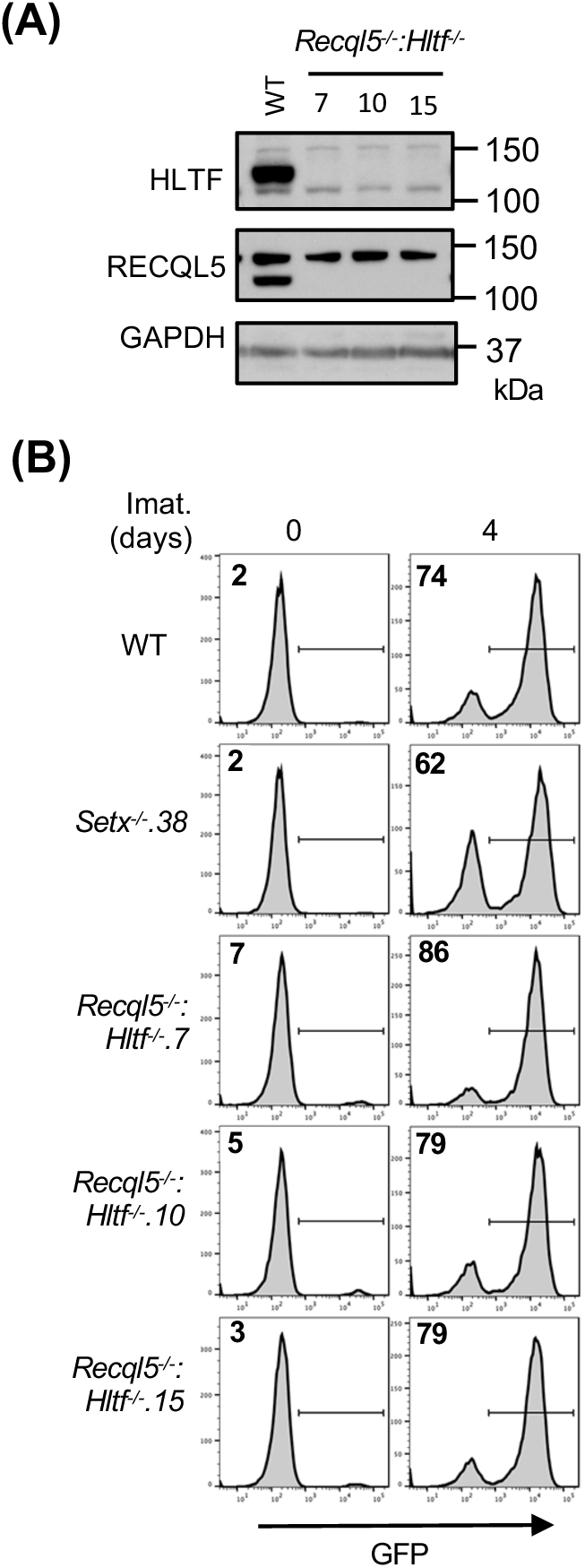
*Recql5*^-/-^*: Hltf*^-/-^ abl pre-B cells exhibit normal V(D)J recombination. (A) Western blot analysis of cell lysates from WT and three independently isolated *Recql5*^-/-^: *Hltf*^-/-^ abl pre-B cells using HLTF, RECQL5 and GAPDH antibodies. (B) Flow cytometric analysis of GFP expression from pMG-INV in WT, *Setx*^-/-^ and three clones of *Recql5^-/-^*: *Hltf^-^*^/-^ abl pre-B treated with imatinib (imat.) for the indicated times. The percentages of GFP^+^ cells are indicated in the top left corners of the histograms.

**Figure S7:**
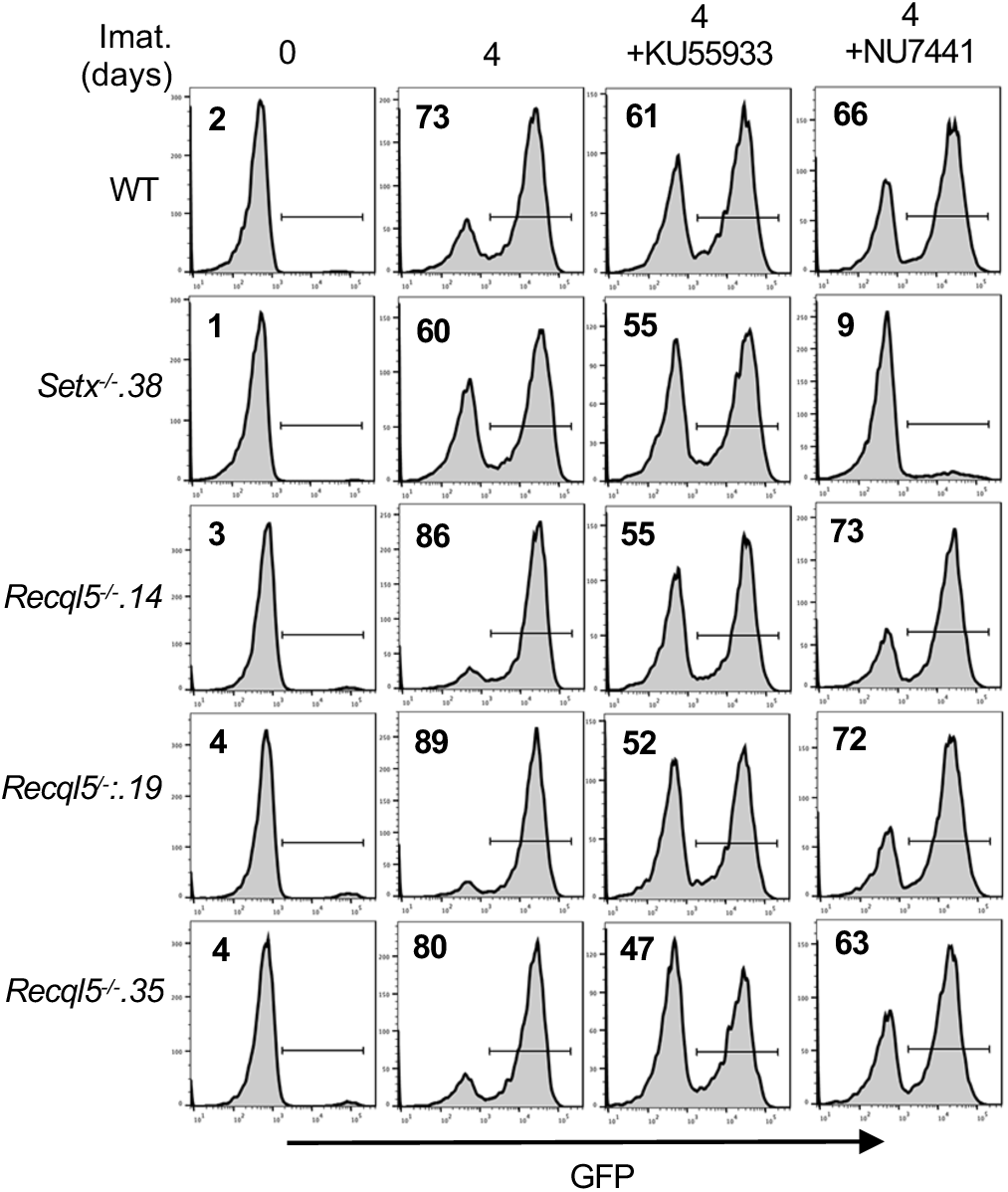
V(D)J recombination in *Recql5*^-/-^ abl pre-B cells. Flow cytometric analysis of GFP expression from pMG-INV in WT, *Setx*^-/-^ and three clones of *Recql5^-/-^* abl pre-B treated with imatinib (imat.) alone or in the presence of the ATM kinase inhibitor KU55933 and the DNA-PKcs kinase inhibitor NU7441 for the indicate times. The percentages of GFP^+^ cells are indicated in the top left corners of the histograms.

**Figure S8:**
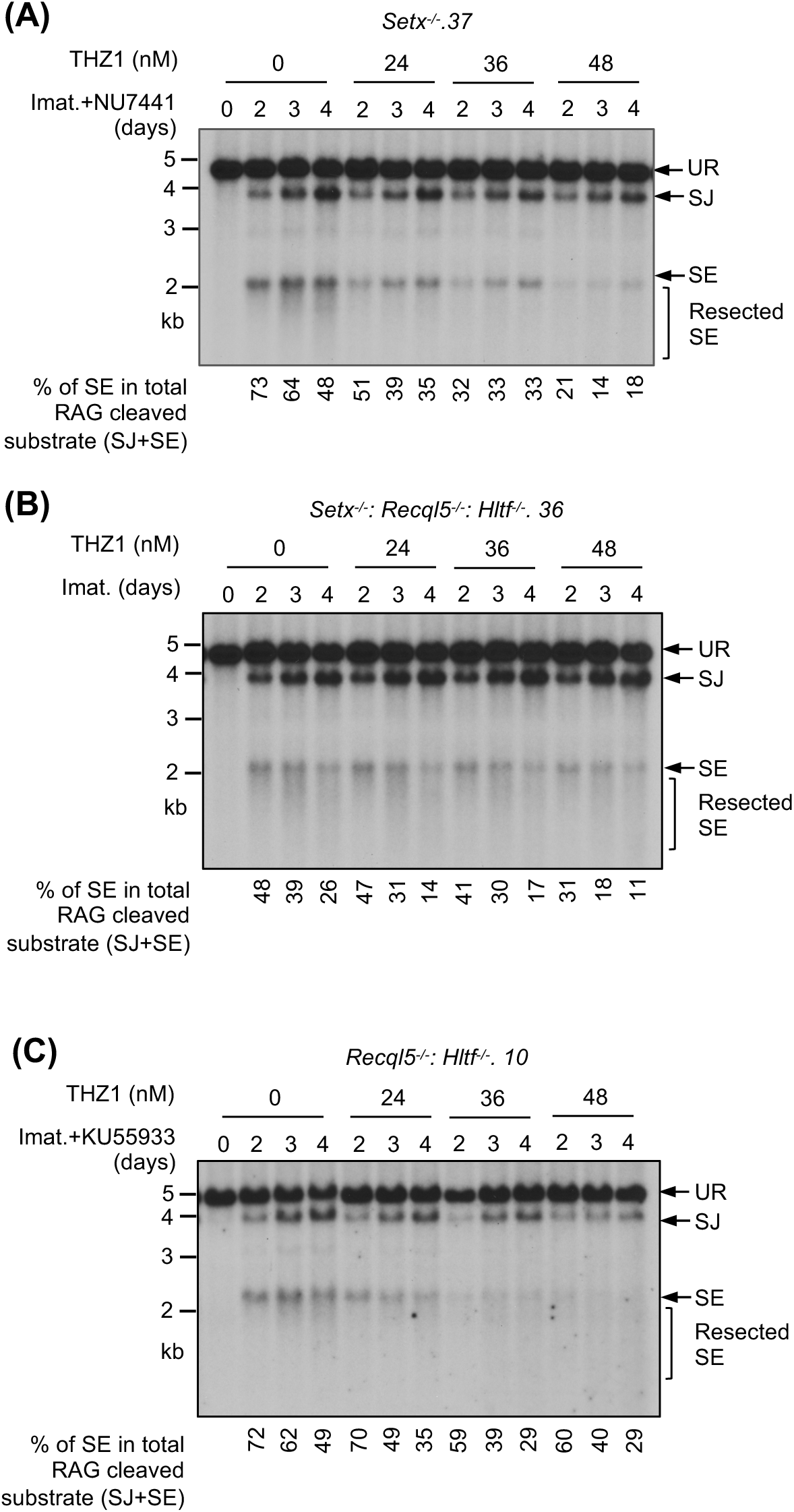
Partial inhibition of RNA polymerase II improves NHEJ-mediated RAG DSB repair in helicase-deficient abl pre-B cells. Southern blot analysis of *EcoR*V-digested genomic DNA from pMX-DEL^SJ^ containing, imatinib (imat.) treated *Setx*^-/-^ (with the DNA-PKcs kinase inhibitor NU7441) (A), *Setx*^-/-^: *Recql5*^-/-^*: Hltf*^-/-^ (B) and *Recql5*^-/-^*: Hltf*^-/-^ (with the ATM kinase inhibitor KU55933) (C) abl pre-B cells with increasing concentrations of the CDK7 inhibitor THZ1 for the indicated times.

## Materials and Methods

### Cell cultures and cell line generation

WT (M63.1: pMG-INV 36: iCas9.302) abl pre-B cells with chromosomally integrated pMG-INV reporter and pCW-Cas9 (Addgene #50661) were described previously (*29*). B.WT abl pre-B cells were immortalized from primary pre-B cells harvested from mice obtained from the Jackson Laboratory (strain#029415) following the same procedure (*29, 69*). To generate clonal abl pre-B cell lines deficient in genes investigated in this work, abl pre-B cells with integrated pCW-Cas9 were treated with 2 μg/ml doxycycline for 2 days to induce Cas9 expression, followed by electroporation with the gRNA-containing lentiviral vector pKLV-hCD2 using the 4D Nucleofector (Lonza) with the SG Cell Line 4D-Nucleofector™ X Kit L and the pulse code CM147. 24 hours after electroporation, gRNA-expressing cells were sorted based on hCD2 expression by MACS cell separators and MS columns (Miltenyi Biotec). Clonal deficient mutant cells were obtained by limiting dilutions and the inactivation of targeted genes in the resulting cell lines were verified by Sanger sequencing (*Setx*^-/-^) or western blot (*Recql5^-/-^*, *Hltf*^-/-^ and *Prkdc*^-/-^). To sequence mutated *Setx* alleles, amplicons containing *Setx* exon 4, where the *gSetx* targets, were generated by PCR and cloned into the pCR-Blunt-II-TOPO vector (Invitrogen) for sequencing.

To generate abl pre-B cell lines with a triple HA (3HA) epitope tag in-frame inserted at the 3’ end of the endogenous *Setx* coding sequence, a recombination donor containing the 3HA coding sequence flanked by ∼800 bp genomic sequences immediately 5’ or 3’ to the stop codon of *Setx* was generated by PCR, cloned into pCR-Blunt-II-TOPO vector (Invitrogen), and verified by sequencing. The sequence of a gRNA targeting near the *Setx* stop codon was included at the 5’ and 3’ end of the recombination donor DNA for liberating the linear donor DNA fragment upon transfection into abl pre-B cells expressing the same gRNA and Cas9. The recombination donor and the gRNA expressing plasmids were transfected to WT abl pre-B cells expressing Cas9 using the 4D Nucleofector (Lonza) with the SG Cell Line 4D-Nucleofector™ X Kit L and the pulse code CM147. The resulting cells, after selected for hCD2 expression from the gRNA expressing vector, were subject to limiting dilution to isolate single cell clones. Clonal *Setx^3HA/3HA^* abl pre-B cells were verified by PCR amplification with primers outside of the *Setx* homology donor sequences and *BamH*I digestion (*BamH*I recognition sequences reside in the 3HA coding sequence) of the resulting amplicons. PCR amplicons from cells with both *Setx* alleles successfully targeted could be completely digested by *BamH*I while amplicons from the WT allele remained resistant to *BamH*I.

Abl pre-B cell lines carrying missense mutations K1945R (*Setx^K1945R/K1945R^*) and L2131W (*Setx^L2131W/L2131W^*) were generated by targeted integration of the recombination donor DNA constructs containing the indicated mutations and 800 bp *Setx* genomic sequences 5’ and 3’ of the mutations in *Setx^3HA/3HA^* abl pre-B cells as described above. Synonymous mutations were incorporated in the targeting constructs to create recognition sequences for *EcoN*I (K1945R targeting construct) and *Xho*I (L2131W targeting construct). Biallelically targeted *Setx^K1945R/K1945R^* and *Setx^L2131W/L2131W^* mutant clones were verified by digesting PCR amplicons covering the respective mutations using primers outside of the donor sequences with *EcoN*I and *Xho*I, respectively.

All cell-lines were grown in DMEM (no L-glutamine, GIBCO, 11960077) supplemented with 10% heat-inactivated fetal bovine serum (GIBCO, A52567-1), 100 U/mL penicillin/ streptomycin (GIBCO, 15140122), 1mM sodium pyruvate (GIBCO, 11360070), 2mM L-glutamine (GIBCO, 25030081), 1X non-essential amino acids (GIBCO, 11-140-050), and 55μM beta-mercaptoethanol (Sigma Aldrich, M3148_100ML) at 37°C. Imatinib (S2475, used at 3μM), ATM kinase inhibitor KU55933 (S1092, used at 15μM), DNA-PKcs kinase inhibitor NU7441 (S2638, used at 5μM) were from Selleckchem. CDK7 inhibitor THZ1 (9002615) was from Cayman Chemical.

### Genome-wide CRISPR/Cas9 screens

Four abl pre-B cell lines with integrated Tet-ON Cas9 transgenes (pCW-Cas9) were used in genome-wide CRISPR/Cas9 gRNA screens – WT (M63.1: pMG-INV 36: iCas9.302), *Setx*^-/-^. 38, *B. Setx*^-/-^.10 and *Setx^-/-^: Recql5^-/-^. 5*. For each screen, ∼180×10^6^ cells were transduced with a lentiviral gRNA library containing 90230 gRNAs targeting 18424 mouse genes (Addgene #67988) at a 40-50% transduction efficiency, as determined by the percentage of BFP+ cells after transduction (*70*). Three days after transduction, stably transduced cells expressing BFP were isolated by fluorescence-activated cell sorting (FACS) and treated with 2μg/ml doxycycline for 6 days to induce Cas9 expression and genome-wide gene inactivation. Cells were then treated with 3μM imatinib for 4 days as described above to induce V(D)J recombination of pMG-INV. Cells that had (GFP^+^) and had not (GFP^−^) undergone pMG-INV recombination were purified by FACS and genomic DNA was isolated from these cells. The gRNAs from each population were amplified using nested PCR and sequenced on an Illumina NextSeq500 platform performed in the Genomic Core Laboratory of Heflin Center for Genomic Sciences at University of Alabama at Birmingham (UAB). Primers used in the PCR reactions are listed in Table S6.

The gRNA sequences were retrieved from FASTQ files using Seqkit (*71*). The derived sequences were mapped to the reference gRNA sequences of the library (*70*). The number of reads of each gRNA were normalized as follows: Normalized reads of a particular 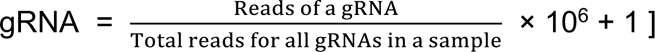 (*72*). The enrichment score of a gRNA is calculated as the ratio of normalized reads of the gRNA between two samples (GFP^−^ vs. GFP^+^).

### Flow cytometric analysis

GFP expression of imatinib-treated abl pre-B cells was acquired on LSRFortessa X-20 analyzers (BD Biosciences) supported by the Flow Cytometry and Single Cell Core and Stem Cell Shared Facility of Division of Hematology and Oncology, UAB. Data analyzed was performed using FlowJo software (FlowJo, LLC).

### Southern blot analysis

To analyze V(D)J recombination products and intermediates of pMX-DEL^CJ^ and pMX-DEL^SJ^ by Southern blot, 8 μg genomic DNA purified from untreated, or imatinib-treated abl pre-B cells was digested with *EcoR*V and resolved in 1.2% TAE gels. Upon transferring DNA to Whatman Nytran SPC membranes (Cytiva, 10416296), restriction fragments corresponding unrearranged reporter (UR), coding join (CJ), coding end (CE), signaling join (SJ) and signaling end (SE) were visualized using a human CD4 gene fragment as the probe as previously described (*23*).

### Western blot analysis

For western blot analysis, protein samples were resolved in 4-12% NuPAGE Bis-Tris gels or 3-8% NuPAGE gels followed by wet transfer to 0.45 μm Protran nitrocellulose membranes (Cytiva, 10600002). The membranes were blocked in 5% non-fat milk for 30 min at room temperature and probed with antibodies as indicated. Antibodies to RECQL5 (12468-2-AP) and MRE11 (10744-1-AP) are from Proteintech. HA antibody (901501) is from Biolegends. DNA-PKcs antibody is from Thermo Fisher Scientific (MS-423-P). HLTF is from Bethyl Laboratories (A300-230A). GAPDH antibody is from Sigma Aldrich (G8795). KAP1 antibody is from Cell Signaling Technology (4123).

### Analysis of structural variants of SJs from pMG-INV by next generation sequencing

To determine structural variants of SJs, genomic DNA from imatinib-treated abl pre-B cells was first subject to PCR to generate amplicons 1.1 kb in size with primers pMG-INV SJ F4 and pMG-INV 3’-3 R flanking the junction of SJs. The 1.1 kb amplicon was used as the template in an additional round of PCR with primers PE.P5_SJ23 and P7_SJ12 located 130 bp to the 5’ and 3’ of the SJ junction to generate a ∼260 bp product with Illumina indexes and adaptors for 150 bp paired end sequencing using miSeq Micro platform (Illunima). Primers used in the PCR reactions are listed in Table S6.

The sequencing results were analyzed, and structural variants were identified as previously described (*73*). Briefly, reads were de-multiplexed based on their index sequences. After removal of adapter sequences, low-quality reads, and trimming reads that were shorter than 20 bp by using cutadapt (v1.3.1), filtered reads were aligned with bowtie2 (version 2.1.0) to the 295-bp pMG-INV SJ reference sequence (Supplemental file 1). With mapped reads in BAM format as input, structural variants were called by using Pindel (*74*). Indels and their microhomology status that span the break point site (the 165-nt position in the target region) with ≥5 supporting reads were extracted, counted, summarized and compared among different conditions (Table S5). P-values (Table S5) were calculated by two-tailed Student’s t-test.

