## Supplementary material for "Senataxin and DNA-PKcs Redundantly Promote Non-Homologous End Joining Repair of DNA Double Strand Breaks During V(D)J Recombination": Table S6

**Table S6: Oligo nucleotides sequences.**

| **Name** | **Sequence** | **Note** |
| --- | --- | --- |
| Primers for CRISPR/Cas9 gRNA screen sequencing | | |
| pKLV lib330F: | AATGGACTATCATATGCTTACCGT | For gRNA library sequencing (1^st^ round PCR) |
| pKLV lib490R: | CCTACCGGTGGATGTGGAATG | For gRNA library sequencing (1^st^ round PCR) |
| PE.P5_pKLV lib195 Fwd | **AATGATACGGCGACCACCGAGATCTACAC**GGCTTTATATATCTTGTGGAAAGGAC | For gRNA library sequencing (**P5 adaptor**) |
| P7 index180 Rev: | **CAAGCAGAAGACGGCATACGAGAT**-index-*GTGACTGGAGTTCAGACGTGTGCTCTTCCGATC*CAGACTGCCTTGGGAAAAGC | For gRNA library sequencing (**P7 adaptor,** *Illumina sequencing primer*) |
| gRNA library seq Read 1 primer | GGCTTTATATATCTTGTGGAAAGGACGAAACACCG |  |
| gRNAs for gene inactivation | | |
| *gSetx* | AAGTGACTTACGAGGCGTA |  |
| *gRecql5* | CGTTGCAGCTCGACTCAGG |  |
| *gHltf* | TCTGTACTGCCACATGAGC |  |
| *gPrkdc* | ATGCGTCTTAGGTGATCGA |  |
| Primers for SJ PCR and sequencing | | |
| pMG-INV SJ F4 | CGGCATCAAGGCGAACTTCA | For SJ sequencing (1^st^ round PCR) |
| pMG-INV 3'-3 R | GTAAAGCATGTGCACCGAGG | For SJ sequencing (1^st^ round PCR) |
| PE.P5_SJ23 | **AATGATACGGCGACCACCGAGATCTACAC**CTGACTTGAATCATGTTGTTTTCCAGACTT | For SJ sequencing (**P5 adaptor**) |
| P7_SJ12 | **CAAGCAGAAGACGGCATACGAGAT-**index-*GTGACTGGAGTTCAGACGTGTGCTCTTCCGATC*CCAAGCGGCTTCGGCCAGTAACGTT | For SJ sequencing (**P7 adaptor,** *Illumina sequencing primer*) |
| SJ seq Read 1 primer | CTGACTTGAATCATGTTGTTTTCCAGACTTCAACT |  |
