## Supplemental file 1 for "Senataxin and DNA-PKcs Redundantly Promote Non-Homologous End Joining Repair of DNA Double Strand Breaks During V(D)J Recombination"

```
LOCUS       Signal_join_miSe         295 bp DNA     linear   UNA 30-JUN-2024
DEFINITION  natural linear DNA
ACCESSION   .
VERSION     .
KEYWORDS    .
SOURCE      natural DNA sequence
  ORGANISM  unspecified
REFERENCE   1  (bases 1 to 295)
  AUTHORS   .
  TITLE     Direct Submission
  JOURNAL   Exported Jun 30, 2024 from SnapGene Viewer 6.2.2
            https://www.snapgene.com
FEATURES             Location/Qualifiers
     source          1..295
                     /mol_type="genomic DNA"
                     /organism="unspecified"
     primer_bind     1..30
                     /label=(Partial) PE.P5_SJ23
     iDNA            complement(127..165)
                     /label=23 RSS
     misc_feature    166..193
                     /label=12 RSS
     primer_bind     complement(271..295)
                     /label=(Partial) P7_SJ12
ORIGIN
        1 ctgacttgaa tcatgttgtt ttccagactt caacttgact atcagccaga aattcagtgg
       61 caaaccccct ccacccatcc ctagtgaggg ttcctagtga gggaggaagg actaactcga
      121 gggtgtggtt tttgtacagc cagacagtgg agtactacca ctgtgcacag tgctacagac
      181 tggaacaaaa acagaccctc gttggccgcc accgatctct cgaggtcgac ggtatcgata
      241 agcttgatat cgaattccgc ccccccccct aacgttactg gccgaagccg cttgg
//
```
